# Interpreting *Cis*-Regulatory Interactions from Large-Scale Deep Neural Networks for Genomics

**DOI:** 10.1101/2023.07.03.547592

**Authors:** Shushan Toneyan, Peter K Koo

## Abstract

The rise of large-scale, sequence-based deep neural networks (DNNs) for predicting gene expression has introduced challenges in their evaluation and interpretation. Current evaluations align DNN predictions with experimental perturbation assays, which provides insights into the generalization capabilities within the studied loci but offers a limited perspective of what drives their predictions. Moreover, existing model explainability tools focus mainly on motif analysis, which becomes complex when interpreting longer sequences. Here we introduce CREME, an *in silico* perturbation toolkit that interrogates large-scale DNNs to uncover rules of gene regulation that it learns. Using CREME, we investigate Enformer, a prominent DNN in gene expression prediction, revealing *cis*-regulatory elements (CREs) that directly enhance or silence target genes. We explore the intricate complexity of higher-order CRE interactions, the relationship between CRE distance from transcription start sites on gene expression, as well as the biochemical features of enhancers and silencers learned by Enformer. Moreover, we demonstrate the flexibility of CREME to efficiently uncover a higher-resolution view of functional sequence elements within CREs. This work demonstrates how CREME can be employed to translate the powerful predictions of large-scale DNNs to study open questions in gene regulation.

## Introduction

Recent advances in sequence-based genomic deep neural networks (DNNs) have shown notable success in predicting gene expression by considering significantly larger inputs^1–4^, which aim to capture the influence of distal *cis*-regulatory elements (CREs). These DNNs bring promise to decode the *cis*-regulatory codes that drive differential gene expression across cell types, predict the effects of genetic variation, and design novel regulatory sequences with desirable properties. However, the extensive sequence size of large-scale DNNs presents a challenge when evaluating their predictions and interpreting learned patterns.

Current methods for evaluating large-scale models have relied on assessing the alignment between predictions and existing experimental perturbation assays^1,5,6^—such as massively parallel reporter assays^7,8^ and CRISPR interference (CRISPRi)^9^—as well as statistical analyses like expression-quantitative trait loci^6,10,11^. However, these only provide a narrow evaluation of how well DNN predictions agree with the specific biological question being probed within the studied loci. Moreover, the underlying biology can be difficult to assess because different experimental technologies introduce distinct biases and noise sources, which do not generalize across experimental technologies.

Conversely, prevailing post hoc model interpretability methods concentrate primarily on the analysis of motifs^12–22^, short DNA sequences associated with regulatory functions. As sequence inputs for DNNs grow longer, deciphering the complex coordination of motifs at a scale of hundreds of kilobases (kb) becomes increasingly difficult to interpret.

To bridge this gap, we present CREME (*Cis*-Regulatory Element Model Explanations), an *in silico* perturbation toolkit designed to examine large-scale DNNs trained on functional genomics data. In contrast to existing model interpretability methods, CREME can provide interpretations at various scales, including a coarse-grained CRE-level view as well as a fine-grained motif-level view. CREME is based on the notion that by fitting experimental data, the DNN essentially approximates the underlying “function” of the experimental assay. Thus, the trained DNN can be treated as a surrogate for the experimental assay, enabling *in silico* “measurements” for any sequence assuming generalization under covariate shifts (i.e., predictions are reliable outside the distribution of training sequences). Drawing inspiration from CRISPRi^23,24^, CREME comprises a suite of multi-scale *in silico* perturbation experiments to uncover interactions between CREs and their target genes learned by DNNs (Fig. 1a). Unlike wet-lab experiments, the perturbations by CREME are performed *in silico*, so there are minimal restrictions on the scale of perturbations that can be applied. Thus, CREME employs controlled experiments that enable calibrated claims of gene regulation rules through the lens of genomic DNNs.

**Figure 1.**
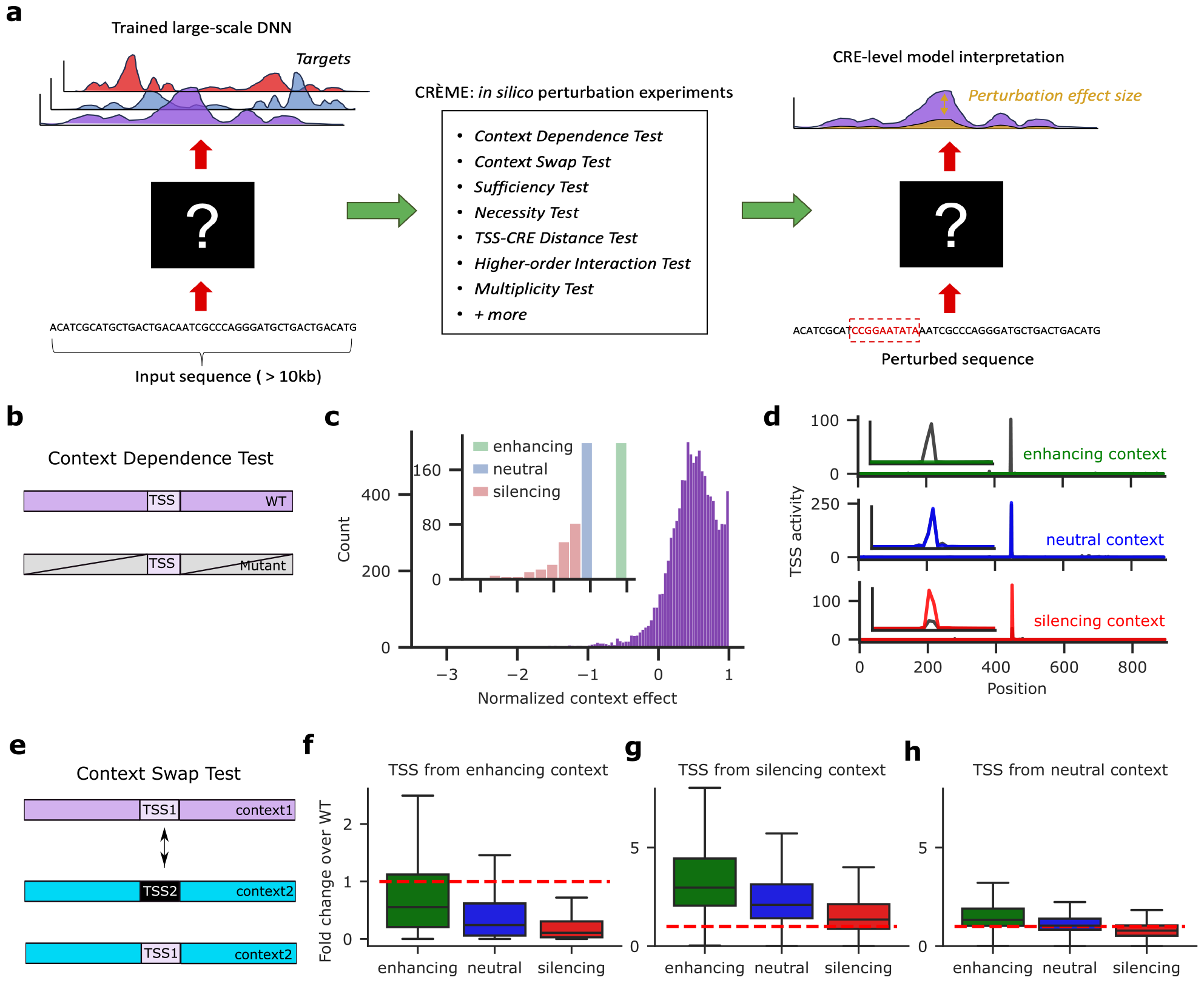
CREME overview and results for context perturbations in K562 cells using Enformer. **a**, CREME offers a suite of *in silico* perturbation experiments that probe specific biological hypotheses. Perturbations are applied to the input sequence and the effect size is measured according to the change in model predictions. **b**, Schematic of Context Dependence Test; the sequence context is perturbed via a dinucleotide-shuffle while keeping the central TSS tile (5 kb) intact. **c**, Histogram of normalized context effect for 10,000 sequences that contain an active, annotated gene in K562 cells. Inset shows the subset of sequences categorized as enhancing, silencing and neutral contexts (N = 200 randomly selected sequences for each context). **d**, Representative sequences from the three context categories showing Enformer’s predictions before and after a context perturbation, with a zoomed in version shown in the inset. **e**, Schematic of Context Swap Test. **f**-**h**, Box-plots of normalized fold change over wild-type (WT) – TSS activity predictions of chimeric divided by wild-type sequences. Results are organized according to the original context categories: enhancing (**f**), silencing (**g**), and neutral (**h**). The number of data points in each box-plot represent an all-vs-all comparison of each respective TSS in each possible context (40,000 data points in each box). Box-plots show the first and third quartiles, the median (central line) and the range of data with outliers removed (whiskers).

To demonstrate the utility of CREME, we interpret Enformer^1^, a DNN that takes *∼*200 kb DNA sequences as input and predicts the corresponding read coverage profiles for 5,313 experiments that includes chromatin accessibility, transcription factor binding, histone marks, and gene expression across various human cell lines and tissues. In this study, we investigate the regulation of gene expression in K562, GM12878, and PC-3 cells. Using a curated list of sequences centered on transcription start site (TSS) annotations from GENCODE^25^ (see Methods), we examine how specific sequence perturbations affect gene expression given by Enformer’s predictions of TSS activity given by CAGE-seq (cap analysis gene expression sequencing) track for the corresponding cell type under study. The results are organized according to specific biological questions.

## Results

### To what extent does distal context affect gene expression prediction?

Gene regulation relies on the complex interactions of distal regulators^26,27^. While Enformer’s predictions have been shown to depend primarily on individual enhancers located near promoters^6^, the extent to which Enformer relies on a broader set of contexts is unclear. To quantify the extent to which distal context affects TSS activity, we developed the *Context Dependence Test* (see Methods). Briefly, this test measures the effect of shuffling the sequence context beyond the proximal regions (*∼*5 kb) centered on the TSS-under-investigation (Fig. 1b). To reduce the effects of spurious patterns, we performed 100 independent dinucleotide shuffles and averaged predictions of TSS activity. We considered the top 10,000 genes with the highest predicted CAGE activity at the TSS for each cell line. The shuffling and averaging of predictions is a procedure similar to global importance analysis (GIA^15^), which serves to marginalize out the effects of the context on TSS activity. As described in the GIA guidelines^15^, this perturbation procedure assumes that the dinucleotide-shuffled sequences are neutral and not out-of-distribution for the model. Previous studies have also employed a similar perturbation using Enformer^6,28^.

The context effect was quantified by calculating the difference in the predicted TSS activity for the wild-type and mutant sequences, normalized by the wild-type TSS activity to control for variable expression levels (see Methods). Under this *normalized context effect* metric, a positive value indicates that shuffling the context decreases TSS activity, with a maximal decrease in TSS activity resulting in a value of 1. On the other hand, negative values in the normalized context effect scores correspond to increasing TSS activity upon shuffling the context.

Enformer exhibited a wide range of interesting responses to the Context Dependence Test. The majority of context shuffles resulted in positive context effects, which presumably arise due to the disruption of enhancers (Figs. 1c and 1d). We also observed cases where TSS activity did not change, yielding a normalized context effect around 0. This suggests the TSS activity was robust across shuffled contexts for these genes, which can arise from a neutral context (i.e., depletion of CREs), the presence of several CREs with a net neutral effect, or TSS activity that is independent of the context. In rarer cases, we observed increasing TSS activity upon shuffling the context, suggesting that active silencing elements were disrupted^29–32^. Based on these results, we organized a subset of the sequences—200 sequences randomly sampled for each of the three context categories (i.e., enhancing, silencing, and neutral)—for further analysis (Fig. 1c, inset).

To test the generalization of the Context Dependence Test, we repeated the same analysis for GM12878 and PC-3 cell lines and observed similar context effect distributions (Extended Data Figs. 1a and 1b). By directly comparing the normalized context effects for the same genes across different cell lines, we observed a high correlation, on average (Extended Data Fig. 1c). However, for certain genes, the normalized context effects exhibited flipped values, i.e., positive effect in one cell line and negative effect in another cell line. This suggests that gene context harbors regulatory elements that are enhancing in one cell type and silencing in other cell types. However, the coarse-grained resolution of the Context Dependence Test cannot specify whether the observed differential behaviors are driven by the same sequence elements or from different CREs throughout the context.

We also employed the Context Dependence Test for Borzoi^4^, an ensemble of four large-scale DNNs that employs a larger input size of 524 kb and predicts strand-specific CAGE outputs (in addition to many other tracks), and observed similar trends across the three cell lines (Extended Data Fig. 2). Due to the size of Borzoi, we limited shuffle perturbations to 5 total shuffles for each of the four models and averaged all sequence predictions across the ensemble. Interestingly, Borzoi’s predictions generally reflect weaker effect sizes for context shuffles; Enformer yielded more values closer to 1 and a higher proportion of negative values. We suspect Enformer might be placing a stronger emphasis on the smaller numbers of CREs that it considers within its smaller receptive field. Since generating predictions with Borzoi is more computationally expensive than Enformer, we continue to investigate Enformer while making comparisons with Borzoi when feasible.

**Figure 2.**
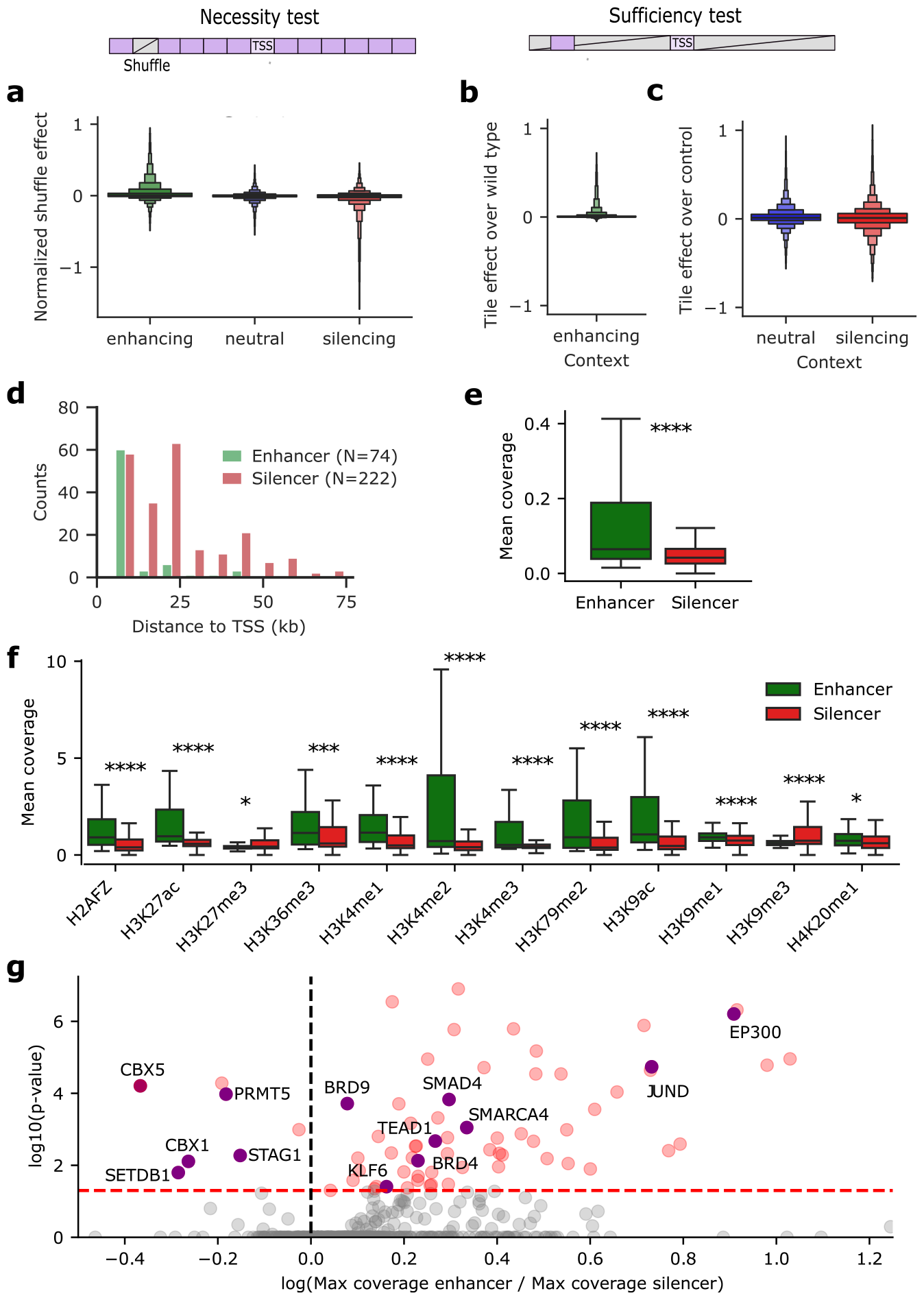
CRE-level analysis in K562 using Enformer. *Top*, Schematic of the Necessity Test and Sufficiency Test. Necessity Test: a TSS-centered sequence with a second shuffled tile indicated by a grey, slashed box. Sufficiency Test: a TSS tile and tile of interest are embedded in dinucleotide-shuffled sequences at their original positions. **a**, Boxen plot of the normalized shuffle effect for each tile in sequences from enhancing, neutral and silencing context categories (Necessity Test). **b, c**, Boxen plot of tile effects normalized by wild-type (**b**) and control, i.e. the intrinsic TSS activity (**c**) for each tile in sequences from enhancing, neutral and silencing context categories (Sufficiency Test). **a**-**c** Boxen-plots have 7,600 context-derived tiles in each sequence context corresponding to 38 tile shuffles in each of the 200 sequences. **d**, Histogram of the distance between CRE tiles from TSS for 74 sufficient enhancer and 222 sufficient silencer tiles, defined by sufficiency thresholds (enhancers > 0.3 from **b** and silencers < -0.3 from **c**). **e**,**f** Boxplots of mean DNase-seq coverage (**e**) and mean histone mark coverage (**f**) of sufficient enhancer and silencer tiles. Number of data points in green and red boxes equal 76 and 222, respectively. Statistical significance is given by the Mann–Whitney U test (*: *p* < 0.05; ***: *p* < 0.001; ****: *p* < 0.0001). Boxplots show the first and third quartiles, the median (central line) and the range of data with outliers removed (whiskers). **g**, Scatter plot of TF enrichment analysis given by the significance level according to the Mann–Whitney U test (adjusted for multiple testing) versus the log-fold enrichment of maximum TF ChIP-seq coverage in sufficient enhancers versus silencers. Some activating and repressive TFs are annotated.

### How compatible are contexts for different genes?

Next, we explored how gene expression changes when a TSS along with its proximal context (5 kb) is inserted into different, non-native (but genomic) contexts using CREME’s *Tile Swap Test* (Fig. 1e). To quantify the effect of these chimeric swaps, we computed the *fold change over wild type* – the ratio of the predicted TSS activity for the mutant sequence and the wild type sequence. We tested the full combination of TSS-context pairs (all-versus-all, comprising 360,000 total sequences in K562) and stratified the results according to the context category of the source TSS tile.

Interestingly, swapping a TSS from an enhancing context into other enhancing contexts resulted in a *∼* 50% decrease in TSS activity, on average (Fig. 1f, middle). This suggests that according to Enformer enhancers can be somewhat effective at enhancing any other gene, but they are better tuned for their native gene, perhaps through some compatibility rules^33–35^. A more considerable drop in TSS activity was observed when placed into non-enhancing contexts, which suggests that, according to Enformer, neutral and silencing contexts are depleted of enhancers or their enhancers have substantially weaker effects.

On the other hand, we found that silencing context acts more generically; silencing context remains effective at silencing any other gene that was originally from silencing context (Fig. 1g). Swapping the TSS from silencing context into non-silencing context led to a substantial increase in TSS activity. This suggests that other contexts, especially enhancing context, are depleted of silencing elements or are outcompeted by enhancers. Interestingly, swapping the TSS from neutral contexts into other contexts resulted in modest changes to the original TSS activity levels (Fig. 1h).

As expected, the genes from neutral context were enriched for housekeeping genes^36^ (22%) compared to the other contexts (5% for enhancing context and 5% for silencing context). The same trends in the Context Swap Test were shared across other cellular contexts (Extended Data Figs. 1d and 1e). Together, our results suggest that Enformer considers context (e.g., enhancing or silencing context) is influential in its prediction of TSS activity across a wide range of genes.

### Which CREs are necessary for gene expression?

The previous analyses provided a coarse-grained view of how distal sequence context (beyond 5 kb) influences TSS activity. Using CREME’s *Necessity Test*, we can map the locations of putative enhancers and silencers within these extended sequence contexts and quantify their effect size on TSS activity (Fig. 2a). Specifically, we tiled the input sequence into non-overlapping 5 kb bins and monitored how TSS activity is influenced by shuffling each tile independently. Henceforth, we performed 10 shuffles per data point or sequence when using Enformer. Importantly, we define a CRE as a 5 kb sequence element that influences TSS activity. The choice of 5 kb tiles enables a higher resolution view of CREs within Enformer’s extended context, encompassing the average perturbation length of experiments probing enhancer function, e.g., CRISPRi^37^, while remaining computationally tractable for the scale of perturbation experiments considering the cost of making predictions using Enformer. CREME can accommodate other choices in tile sizes; later, we explore smaller tile sizes (down to 50 bp) with an efficient search strategy.

We performed the Necessity Test on every context tile in all sequences from each context category (N = 22,800 total tile perturbations). The effect size was quantified by calculating the difference in predicted TSS activity between the wild-type and mutant sequences, followed by a normalization with the predicted wild-type TSS activity to make relative comparisons across genes (see Methods). Under this *normalized shuffle effect* metric, a positive value indicates an enhancing effect, negative values indicate a silencing effect, and a value of 0 indicates no effect.

We found that individual tiles in enhancing backgrounds tend to influence TSS activity positively, on average. In contrast, tiles in silencing context are enriched to yield a negative influence (Fig. 2a). Notably, all contexts, including neutral context, appear to display a mix of CRE behaviors, such as enhancers and silencers, with most tiles having little to no effect on TSS activity. We observed the same trends across the different cell lines (Extended Data Figs. 3a and 3b). This suggests that according to Enformer, sequence elements at 5 kb resolution exhibit a continuous range of enhancing and silencing effects on gene expression, most of which contribute weakly.

**Figure 3.**
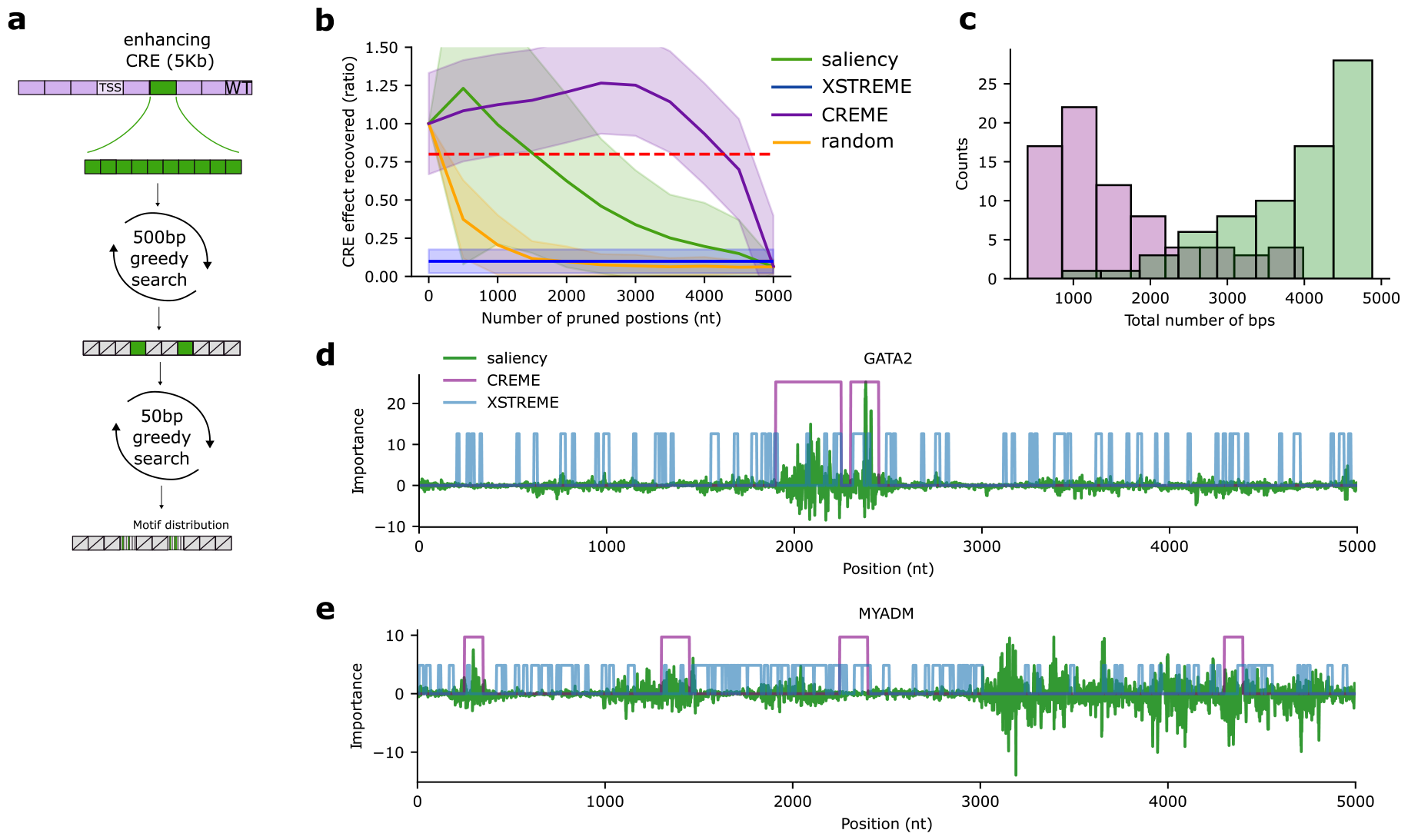
Fine-Tile Search results for enhancing tiles in K562. **a**, Schematic of the Fine-Tile Search procedure. Green 5 kb tile represents a previously identified sufficient enhancer tile. Greedy search aims to prune the least enhancing sub-tiles, first at a 500 bp sub-tile resolution, followed by a second greedy search at a 50 bp sub-sub-tile resolution among the surviving 500 bp sub-tiles. The final output is a sequence with the core 50 bp elements among randomized sequences that recapitulates the sufficiency of the whole tile. **b**, Plot of the average CRE effect recovered ratio (which is the TSS activity when only sub-tiles are embedded in shuffled context divided by the the activity when the full tile is embedded) versus the number of pruned positions for 74 enhancer tiles using different methods. The red dotted line shows an arbitrary threshold of 80% explained. **c**, Histogram of number of remaining positions at score of 0.8 for CREME versus saliency analysis. **d**,**e** Example annotations of important sequence elements for a putative enhancer for different genes. Note that the importance scale (*y*-axis) is set according to Saliency maps and the heights of binary annotations given by XSTREME and CREME are set arbitrarily for comparison.

### Do individual CREs sufficiently explain TSS activity predictions?

While the necessity test identifies individual CRE tiles needed for TSS activity, it does not specify whether they are sufficient to explain the observed TSS activity levels^38^. To address this, we employed CREME’s *Sufficiency Test*, which embeds a tile-of-interest along with the TSS tile into dinucleotide-shuffled sequences at their original positions and predicts their TSS activity (Fig. 2b; N = 22,800 total tile perturbations). This GIA experiment uncovers the global importance of the combined tiles while effectively removing contributions from background context^15^.

Due to differences in the intrinsic TSS activity levels across contexts, we used separate metrics to provide an intuitive characterization for sequences categorized as enhancing context and the other contexts (see Methods). For enhancing context, we quantified the *tile effect over wild type*, which is the difference in the predicted TSS activity between the context-shuffled sequences with both TSS and tile embedded from just the TSS alone divided by the predicted wild type activity. Under this metric, a normalized effect of 1 indicates that the context tile fully recovers the wild-type TSS activity. For silencing and neutral contexts, we used the *tile effect over control*, which uses a different divisor – intrinsic TSS activity levels (i.e., the context-shuffled sequence with just the TSS), due to the higher activities upon context shuffles in silencing context. Under this metric, a value of 0 indicates that the tile does not change the TSS activity, a value of 1 indicates that the tile doubles the TSS activity, while negative values reflect a decrease in TSS activity.

In all three cell lines, we found that individual tiles in enhancing context behave strictly as enhancers or have no effect (Figs. 2b, Extended Data Figs. 3c and 3d). Most tiles exhibited minor positive effects, with very few tiles driving substantial TSS activity approaching wild-type levels. On the other hand, individual tiles in a silencing context contain strong enhancers and silencers relative to the intrinsic TSS activity levels (Fig. 2c, Extended Data Figs. 3c and 3d). Hence, the observed net silencing effect of the entire context suggests that each context is comprised of an imbalance of silencer tiles compared to enhancing tiles, or the fewer silencer tiles interact synergistically in a non-additive manner, thereby overpowering the strong enhancers. In a neutral context, individual tiles contain a balanced set of weak enhancers and silencers, which cancel out each other’s effect on TSS activity (Fig. 2c, Extended Data Figs. 3c and 3d). Evidently, the necessity of a tile does not imply its sufficiency (Extended Data Fig. 3e).

By performing the Sufficiency Test using Borzoi – based on the enhancing, silencing, and neutral contexts defined by Borzoi’s results for the Context Dependence Test (Extended Data Fig. 2) – we observed similar trends (Supplementary Fig. 1). Notably, there was a mix of enhancing and silencing elements in all three cell lines, albeit with a narrower range of effect sizes compared to Enformer.

### Characterization of sufficient CREs

Next, we evaluated the properties of sufficient tiles that behave as strong enhancers or silencers. Specifically, we identified strong enhancers and silencers according to a stringent threshold based on the sufficiency test, leading to 74 enhancing and 222 silencing tiles (see Methods). We first explored the distance distribution of these putative CREs with respect to their target TSS. Our findings in all three cell lines indicate that the majority of strong enhancer tiles recognized by Enformer are located in close proximity to their target TSS, and the number of enhancers progressively diminishes as the distance increases (Fig. 2d, Extended Data Fig. 4a), in agreement with previous observations^6^. Interestingly, the observed silencer tiles recognized by Enformer exhibit a broader distribution compared to enhancer tiles but are still largely located nearby the target TSS (Fig. 2d, Extended Data Fig. 4a).

**Figure 4.**
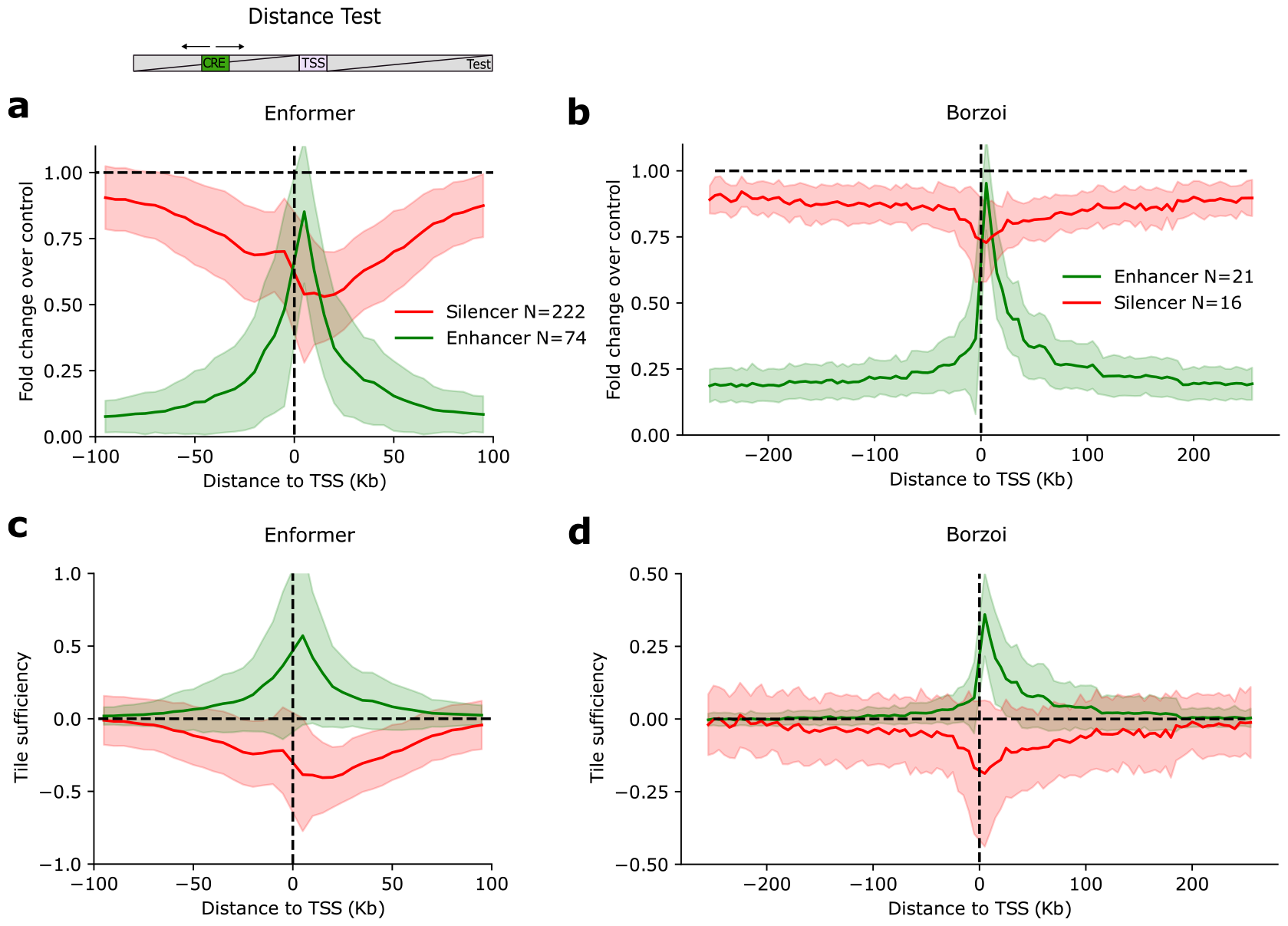
TSS-CRE Distance Test schematic and results. *Top*, Schematic of Distance Test. **a**,**b**, Average plot of the fold change over max of moving a CRE tile along different positions in a sequence (5 kb steps) about a fixed TSS tile in shuffled sequences; max represents the maximum TSS activity across all embedded positions for Enformer (**a**) and Borzoi (**b**). Shaded region represents standard deviation of the mean. **c, d**, Plot of the tile sufficiency versus distance to TSS. Tile sufficiency is calculated according to the predicted TSS activity with a TSS-CRE pair at a given distance minus the control sequence (shuffled context with just the TSS) divided by the WT sequence for enhancers and by the control sequence for silencers.

Next, we compared epigenetic features associated with enhancer and silencer tiles using ENCODE^39^ tracks for each respective cell line. In particular, we averaged the read coverage for available histone marks, chromatin accessibility, and TF ChIP-seq tracks for the tile under study (see Supplementary Data 1 for a complete list of tracks for each cell type). As expected, enhancer tiles were enriched for higher chromatin accessibility, H3K27ac, and H3K9ac, among other histone marks that are well-recognized features of enhancers^40^ (Figs. 2e and 2f). On the other hand, silencer tiles had lower levels of chromatin accessibility and were enriched for H3K9me3 (Figs. 2e and 2f), which are associated with repressing gene expression^41^. These observations were shared across all tested cell types, with the major difference being H3K36me3 and H3K4me3 marks, which were enriched differentially across cell types (Extended Data Figs. 4b-4d).

Using RepeatMasker^42^, we found that silencer tiles had a relatively higher percentage of long interspersed nuclear elements (LINEs, 16.6% versus 9.7%) and long terminal repeats (LTRs, 7.4% versus 2.6%) compared to enhancer tiles – an observation that held across all three tested cell lines (Supplementary Table 1). Similarly, we checked whether the sufficient enhancer and silencer tiles overlapped with annotations for promoters (using GENCODE annotations^25^) or enhancers (using an enhancer atlas for K562^43^). We observed that 27% of enhancer tiles and 24% of silencer tiles overlap with these annotations (Supplementary Table 4). Hence, the CREME-identified CREs learned by Enformer mostly do not overlap with existing CRE annotations.

Finally, we utilized the rich repertoire of TF and chromatin remodeler ChIP-seq ENCODE tracks available for K562. Specifically, we compared each TF’s maximum activity (according to read coverage) in putative silencer and enhancer tiles. We found that the CREs identified by CREME correspond well with known biochemical properties of enhancers and silencers (Fig. 2g). Silencers were enriched for known repressor TFs, including CBX5 (chromobox five protein)^44^, PRMT5 (protein arginine methyltransferase 5)^45^, SETDB1 (Histone-lysine N-methyltransferase)^46^ within silencers. In contrast, enhancers were enriched for activator TFs and remodellers, including EP300 (histone acetyltransferase)^47^, BRD9 (Bromodomain-containing protein 9)^48^, SMARCA4 (SWI/SNF Related, Matrix Associated, Actin Dependent Regulator Of Chromatin, Subfamily A, Member 4)^49^.

Enformer was trained to predict an imbalanced amount of TF tracks in K562 compared to other cell types like PC-3. However, we found that Enformer could still learn similar characteristic features of histone marks and chromatin accessibility across enhancer and silencer tiles across all cell types. This suggests that Enformer can characterize enhancers and silencers from sequence features *de novo*.

Together, these results support that the putative CREs identified by CREME correspond well to the characteristic properties of enhancers and silencers. Unlike traditional approaches, which define CREs according to observational statistics based on biochemical features^50^, CREME identifies putative CREs that directly impact predictions for gene expression in an unbiased manner according to a large-scale DNN.

### Fine-mapping sub-sequence elements that reproduce full CRE effects

CREME tests can be applied at any length scale, but we chose a 5 kb resolution to balance the computational costs of generating Enformer predictions with potential biological insights that could be gained; choosing smaller tile sizes would dramatically increase the number of predictions necessary to carry out each test. Here, we show how CREME can be used to efficiently gain a higher-resolution view of sequence elements that are important within the 5 kb enhancer tiles (N = 74), using a nested greedy search algorithm we call Fine-Tile Search (Fig. 3a). The greedy search aims to identify the set of sub-tiles that can explain the behavior of the whole tile. Here, we evaluate success by calculating the *CRE effect recovered*, the ratio of predicted TSS activity using the sub-tiles versus the whole tile embedded in shuffled sequences. This is equivalent to computing the ratio of sufficiency test for the sub-tiles versus the whole tile. CRE effect recovered of 1 indicates that the set of sub-tiles fully recapitulates the effect of the whole 5 kb enhancer tile.

In each iteration of the greedy search, we pruned non-important tiles by systematically shuffling each tile ten times and averaging the predictions. We then fixed the shuffle for the tile that led to the smallest decrease in the CRE effect recovered and then start another round of the greedy search, focusing on the remaining tiles (see Methods). We deployed the greedy search at two length scales to improve the search efficiency. First, we ran the greedy search to prune 500 bp-sized sub-tiles until a 90% threshold of the CRE effect recovered was reached. We then shifted to a finer scale of 50 bp tiles, searching within the surviving 500 bp tiles. At this resolution our selection criterion relaxed; choosing increments of ten tiles that led to the smallest independent decrease considerably sped up the search. At a 50 bp resolution, the greedy search-stopping criterion was set to 70% of the 5 kb CRE effect recovered.

As a baseline for comparison, we compared CREME’s Fine-Tile Search to a traditional motif enrichment tool called XSTREME^51^ and a standard *post hoc* interpretability method called Saliency Maps^12^, which quantifies nucleotide-level sensitivity of each nucleotide on model predictions. XSTREME analyzed the set of enhancing tiles using known motifs from JASPAR^52^ and identifying *de novo* motifs, providing footprints of all motif hits. Saliency Maps were generated by calculating the gradient of the TSS prediction for a given cell line with respect to the inputs. In each case, we tested how well these methods can identify sufficient sets of positions that recover the whole enhancer tile’s predicted effect on TSS activity. For XSTREME, we used the motif annotations to select sequence elements to be considered for computing CRE effect recovered. For Saliency maps, we set various thresholds to identify different numbers of the most “salient” positions and calculated their CRE effect recovered score.

Strikingly, CREME’s Fine-Tile Search identified more compact subsets of sequences that explained 80% (arbitrary threshold) of the original 5 kb tile compared to saliency-based search and motif analysis (Figs. 3b and 3c). This suggests that saliency maps identify some essential positions but also attribute importance to random nucleotides. Observational motif analysis leads to poor characterization of the sufficient elements within the enhancer tile. Similar trends were shared across different cell types (Supplementary Fig. 2).

In a qualitative comparison, we overlaid the sequence elements identified by each method for putative enhancer tiles (identified by CREME) for GATA2 and MYADM. We observed that saliency maps and CREME agreed on occasion, as seen in the putative enhancer tile for GATA2 (Fig. 3d), but can be quite divergent in other loci, such as the putative enhancer tile for MYADM (Fig. 3e). Importantly, motif hits were rampant throughout each 5 kb locus, with most of the hits not representing sequence elements that drove Enformer’s predicted TSS activity. We observed similar findings across other loci (Supplementary Fig. 3).

Together, the multi-scale Fine-Tile Search is a powerful way to fine-map sequence elements within putative CREs. Unlike attribution methods, which identify important positions either through gradients or via additive approximations^13,14^, CREME identifies a compact set of nucleotides that are necessary and sufficient for model predictions. Due to the computational costs of running Fine-Tile Search experiments, we resume our study defining putative CRE elements at a 5 kb resolution.

### How does a CRE’s effect on a target gene depend on distance?

While the observed enhancer tiles have been found to span a long range of distances to their target TSS (Fig. 2d), it is unclear how their effect size varies with distance to the target TSS. To address this question, we developed the *TSS-CRE Distance Test*, which performs a GIA experiment where the TSS activity is monitored while the distance of a putative enhancer or silencer tile is systematically varied from the target TSS (Fig. 4a). To focus on the trends of the distance-dependent relationship across different genes, we normalized the distance-dependent effect of TSS-CRE pairs by the position of the CRE that yielded the maximum TSS activity (see Methods). A value close to 1 indicates the location of the highest activity for enhancers and the lowest for silencers.

We found that Enformer learns a strong asymmetric distance-dependent relationship of an enhancer tile’s effect on TSS, on average (Fig. 4a). We tested this across 38 positions for 74 enhancer tiles, yielding 2,812 different mutant sequences. This reveals that there is a preferential positioning of the enhancer tile with respect to the TSS; in this direction, the effect size is greatest when juxtaposed next to the TSS tile, with a strong decay with distance, which is similar to trends observed in experimental studies^53–55^. This suggests that, according to Enformer, a weak enhancer can increase its effect on gene expression by moving closer to the TSS in the preferred direction.

By performing the same analysis for TSS-silencer pairs (N = 8,436 mutants), we found a similar trend that shows an asymmetric effect of silencer tiles that decreases with distance from the TSS (Fig. 4a), albeit the effect drops off less rapidly compared to enhancers (Fig. 4b). These distance-dependent relationships were similar across other cellular contexts (Extended Data Fig. 5). While previous studies have suggested that Enformer does not use the full range of its receptive field when modeling distal regulatory elements^6^, our results demonstrate that this mainly pertains to enhancer tiles; Enformer appears to consider the full range of its receptive field for distal silencer tiles.

**Figure 5.**
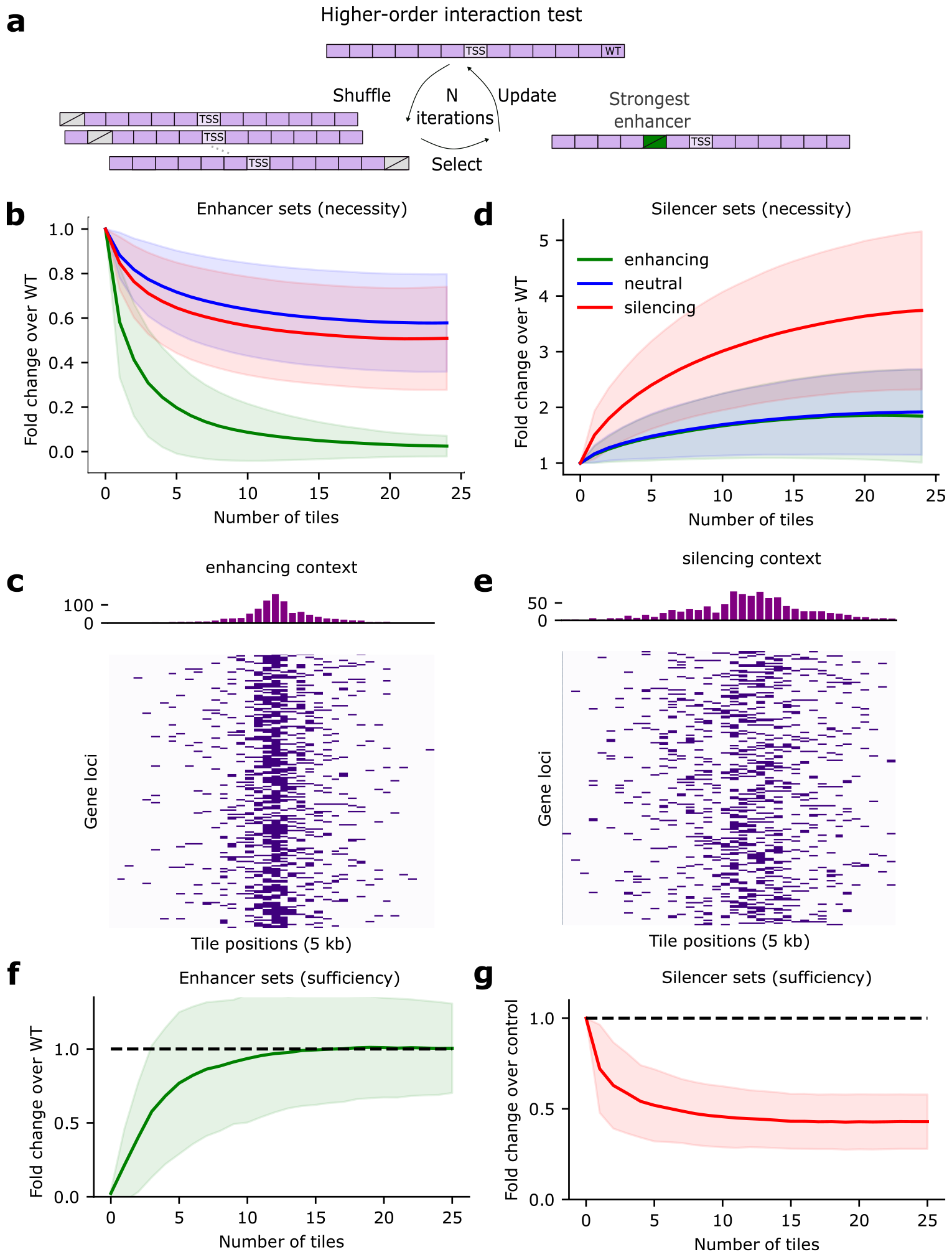
Optimal CRE sets reveal complex interactions for K562 using Enformer. **a**, Schematic of the Higher-order Interaction Test for enhancer CRE sets (grey, slashed boxes represent a shuffled tile and green box indicates an enhancer tile). **b**,**d** Average plot of the greedy search results for enhancer tile sets (**d**) and silencer tile sets (**d**) for sequences from different context categories (N=200 sequences for each context category). The fold change over wild-type (WT) is the predicted TSS activity of the shuffled CRE tiles in each round of the greedy search (indicated by the number of tiles). **c**,**e**, Heatmaps of the locations of the first 5 tiles identified in the enhancer or silencer greedy search within sequences from an enhancing or silencing context, respectively. The histograms on top show the distribution of tile positions. **f**,**g**, Sufficiency of the tile sets identified in each round of greedy search. Average fold change over WT (**f**) and control (**g**), which represents shuffled sequences with just the TSS tile. Sufficiency places the tile sets along with the TSS tile into shuffled sequences. Shaded region represents the standard deviation of the mean.

To test whether these trends are shared with other large-scale DNNs, we repeated the TSS-CRE Distance Test for Borzoi^4^, using enhancer and silencer tiles identified in Borzoi’s Sufficiency Test (Supplementary Fig. 1). For Borzoi, we tested 102 non-overlapping tile positions for the 21 enhancers and 16 silencers yielding 2,142 and 1,632 mutants from enhancers and silencers. Similarly, Borzoi-derived enhancer tiles exhibit a strong asymmetric distance-dependent decay from the TSS, albeit Borzoi appears to have improved upon Enformer by considering more distal enhancers at 100 kb from the TSS (Fig. 4c). On the other hand, the effects of Borzoi-derived silencer tiles were weaker overall and decayed slower with distance relative to Enformer (Fig. 4d). Although the TSS-CRE Distance Test does not probe the exact role of CREs in their native genomic context, this interventional test helps to reveal biases learned by a DNN, mapping out the effective range that CREs affect TSS activity globally, i.e., across sequence contexts.

### How can we identify minimal sets of CREs that regulates gene expression?

To identify minimal sets of CREs that maximally alter TSS activity, we move from single-tile perturbations to multi-tile perturbations using CREME’s *Higher-order Interaction Test* (Fig. 5a). In anticipation that CRE interactions are complex^26,56^, we elected to search for tile sets via an iterative greedy search instead of grouping tiles based on their individual effects. In each round, the greedy search identifies a new tile that yields the largest effect size given the set of tiles found in previous rounds (see Methods); optimizing for higher or lower TSS activity yields sets of silencers or enhancers, respectively. We quantified the effect of multi-tile perturbations with the *fold change over wild type* – normalizing the predicted TSS activity for the mutant sequence by the WT sequence (N = 390,000 combined number of sequences across different contexts in each CRE type). Thus, a greedy search for sets of enhancers/silencers will observe a decrease/increase in the fold change over wild type, respectively. Both tests ultimately converge to TSS activity levels in fully random contexts, that is, the results from the Context Dependence Test.

When probing for enhancing tile sets, we observed that in all cell lines, five enhancer tiles (on average) drive more than 80% of TSS activity for sequences in enhancing contexts (Figs. 5b, Extended Data Fig. 6a). These first 5 CREs are mostly centered around the TSS; however, on an individual sequence basis, CREs can be found to span the entire length of Enformer’s receptive field (Fig. 5c, Supplementary Fig. 4). In contrast, when probing for silencer tile sets, we found that silencing context is enriched for larger numbers of silencer tiles, each with a seemingly smaller effect size (Fig. 5d, Extended Data Fig. 6b). Moreover, the sets of the most effective silencer tiles are distributed more broadly compared to enhancer tiles (Fig. 5e, Supplementary Fig. 5). Interestingly, all contexts, including neutral context, contain enhancer and silencer tiles, albeit with modest effect sizes. Together, this suggests that, according to Enformer, sequence context contains numerous enhancers and silencers and their overall net effect drives the predicted TSS activity.

**Figure 6.**
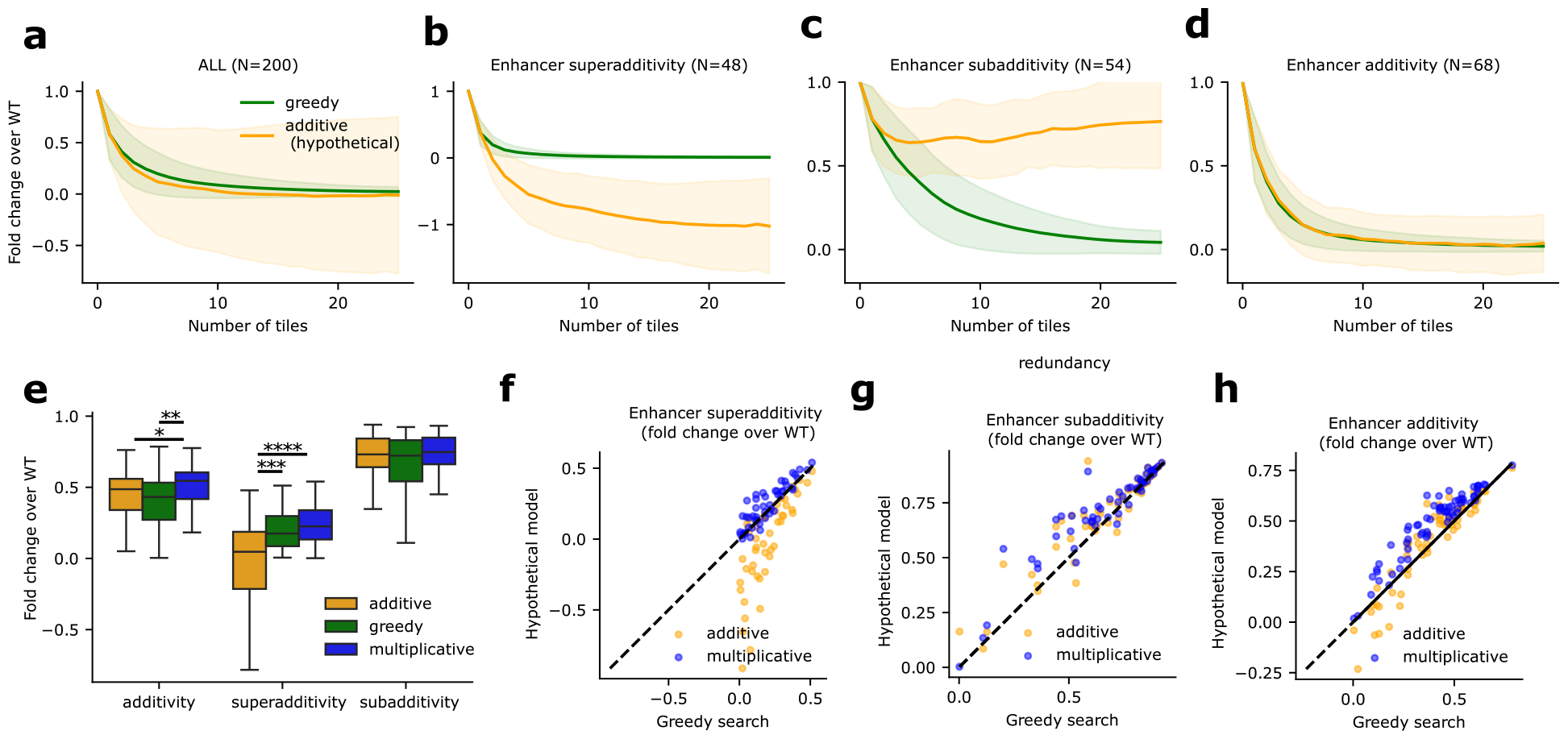
Investigation of CRE interactions. **a-d**, Comparison of the average fold change over control for enhancer sets for sequences categorized as enhancing context versus a hypothetical additive effects model. The 200 sequences from enhancing contexts are stratified according to interaction type, superadditivity (**b**), subadditivity (**c**), and additivity (**d**) using mean squared error based thresholds of 0.1 for superadditivity and subadditivity and 0.05 for additivity definition (with some ambiguous cases left out of classification). Shaded region represents standard deviation of the mean. **e**, Comparison of hypothetical additive model and hypothetical multiplicative model versus greedy search outcomes at iteration 2 of the higher-order interaction test. The number of points in each box is 68, 48 and 54 for additivity, superadditivity, and subadditivity. Statistical significance was given according to the Mann-Whitney U test (*: *p* < 0.05; **: *p* < 0.01; ***: *p* < 0.001; ****: *p*< 0.0001). **f-g**, greedy search versus hypothetical additive or multiplicative models. Scatter plots show a more detailed view of the data in **e** with x-axes showing the higher-order interaction test outcomes and the y-axes showing the hypothetical model outputs (additive or multiplicative). Boxplots show the first and third quartiles, the median (central line) and the range of data with outliers removed (whiskers).

In a complementary set of experiments, we asked whether the sets of enhancers and silencers found from the greedy search are sufficient to explain the TSS activity compared to the whole context (see Methods). Specifically, we inserted the tile sets identified at each round of the Higher-Order Interaction Test and the TSS tile into shuffled background sequences. For enhancing context, we used the *fold change over wild type*, which compares the TSS activity predictions for the mutant sequences with the wild type sequence. For silencing context, we used the *fold change over control*, which compares the TSS activity of the mutant sequence with the randomized sequences with just the TSS embedded, which serves to measure how much the intrinsic TSS activity drops when silencer tiles are added.

For enhancer sets, we observed that five tiles are sufficient to explain 80% of the whole context, but 11 tiles are needed to fully explain the wild-type TSS activity (Fig. 5f, Extended Data Fig. 6c). On the other hand, for silencer sets, only three silencer tiles are needed to suppress 50% of the intrinsic TSS activity; additional tiles reach saturation within round 5-6 (Fig. 5g, Extended Data Fig. 6d). The discrepancy between the necessity-based Higher-order Interaction Test and the sufficiency control experiments for silencer tile sets suggests that there is a floor on how much gene expression can be silenced. Together, these results highlight how CREME can be used to identify sets of CREs that are necessary and sufficient to explain model predictions of TSS activity.

### Do sets of CREs interact in an additive or non-additive manner?

While the Higher-Order Interaction Test can identify sets of CREs, it does not inform how these CREs interact. To gain insights into CRE interactions, we compared the trajectories from the greedy search with a hypothetical scenario where the overall influence of multi-tile perturbations on TSS activity follows an additive effects model. First, we used the independent effects of the tiles quantified via the first round of the enhancer greedy search, adding up the same tile sets found in each round. If the additive assumption were to hold, we expect the effect size to change monotonically with the same tile order chosen by the enhancer greedy search. However, we observed very erratic individual effect sizes (Extended Data Fig. 7). This suggests that tiles exhibit strong dependencies; a perturbation to one CRE modifies the effect size of other CREs.

Next, we directly compared the additive effects model following the same tile sets defined in each round of the greedy search (see Methods). According to Enformer, sets of putative enhancer tiles are largely additive on average (Fig. 6a, Extended Data Figs. 8a and 8b), which is in agreement with previous studies that experimentally examined how enhancers combine^57^.

However, when stratifying the results (see Methods), we observed that enhancers exhibit a range of complex non-additive behaviors (Figs. 6b-6d, Extended Data Figs. 8a and 8b), including superadditivity and subadditivity. Subadditivity refers to when multiple enhancers appear to each have a small effect size on TSS activity when perturbed individually, which can arise due to the presence of other redundant enhancers^58–61^. Superadditivity refers to when two or more CREs depend on each other, which can arise due to synergistic effects like cooperativity^26,56^ or multiplicative effects^6,34,62,63^. For instance, in the case of cooperativity, a perturbation to an individual CRE cooperating with another CRE will result in a considerable decrease in TSS activity as both are necessary. In this case, the hypothetical additive effect will appear stronger than the observed effect when both CREs are perturbed, leading to negative TSS activities (Fig. 6b).

To assess whether the observed superadditivity can be explained by a multiplicative effects model, we repeated the same analysis as the hypothetical additive case, but this time taking the log of the TSS activity (see Methods). Initially, we compared the results for two enhancer tiles (i.e., round 2 of the greedy search). Using the same stratification defined by the additive model, we directly compared the range of effect sizes from each hypothetical model (i.e., additive and multiplicative) to the effect sizes from the greedy search. We found that the hypothetical additive model explained the additive cases well (Figs. 6e and 6h, Extended Data Figs. 8c-8f), while hypothetical multiplicative model explains superadditive cases better (Figs. 6e and 6f). However, this was only the case for the first 2-3 enhancer tile sets, after which the predictions from the hypothetical multiplicative model diverged from the enhancer greedy search (Supplementary Figs. 6 and 7), which suggests synergistic interactions^64^. We also compared multi-tile perturbations with a hypothetical additive model for silencer sets and found that silencers predominantly exhibit superadditivity (Extended Data Fig. 9).

Notably, previous studies have found that Enformer learns a multiplicative model for a single enhancer-promoter interaction^6^. In this study, we find Enformer learns a wide diversity of enhancer-enhancer interaction rules that varies from locus to locus. A more in-depth analysis into the motif-level rules that govern these differential interactions could be an interesting future direction to explore using CREME’s Fine-Tile Search.

### How does Enformer respond to dosage of CREs?

CRE subadditivity can arise if TSS activity follows a sigmoidal function, where saturation has been reached^65–70^. To test whether Enformer’s dosage response to CREs leads to saturation behaviors, we used CREME’s *Multiplicity Test* to measure the effect of greedily inserting multiple enhancers (or silencers) within dinucleotide-shuffled sequences to maximize (or minimize) TSS activity (Extended Data Fig. 10). Here, each round of a greedy search places an enhancer tile in the location that maximally increases TSS activity. Then, that enhancer tile is fixed, and another round places the same enhancer tile among the remaining positions to identify the next location that maximizes the TSS activity. By the end of 15 rounds, the same enhancer tile will be placed in 15 distinct locations. Due to our interest in the shape of the dosage response and not necessarily the magnitude of the response, we opted to normalize the TSS activity according to when CRE is in its original position within control sequences, i.e., dinuc-shuffled sequences (see Methods).

We observed that Enformer’s predictions of gene expression saturate as the dosage of enhancers (or silencers) increases, albeit different genes plateau at different TSS activity levels (Extended Data Fig. 10). This demonstrates that Enformer has indeed learned a non-linear dosage response to CREs (i.e., enhancers and silencers), where each gene’s TSS activity eventually can reach saturation. Thus, the subadditivity cases observed for sets of enhancers likely reflect that the gene’s saturation levels have been reached. Although silencer sets can reach saturation levels as well, we do not observe any cases of subadditivity (Extended Data Fig. 9). This may be due to the biased selection of active genes in K562 used in this study. Further investigation is needed to understand what factors determine the saturation properties, such as why the magnitude of the response varies across genes.

## Discussion

In summary, CREME provides a suite of *in silico* experiments for the unbiased interpretations of large-scale sequence-based DNNs. CREME enables moving beyond the limitations of existing model interpretability methods geared toward motif analysis by focusing on a CRE-level analysis. CREME reveals the rules of gene regulation – such as the dependence of TSS activity on distal context and the complex coordination of CREs – learned by genomic DNNs.

By interpreting Enformer, we found that each sequence context contains numerous enhancers and silencers that interact in complex ways, including additivity, superadditivity (multiplicative and synergistic), and subadditivity, which arises from saturation of the gene’s TSS activity levels – a phenomenon that can be achieved for any gene in enhancing and silencing context. The overall net effect of these complex interactions leads to the predicted TSS activity levels. According to Enformer, we also find that CREs exhibit a continuum of positive and negative effect sizes for putative enhancers and silencers, respectively. This underscores the difficulty in defining enhancers and silencers through arbitrary cutoffs of activity level.

### Limitations

A major limitation can arise when a DNN’s understanding of gene regulation is not aligned with biological reality. In this case, the results of *in silico* experiments by CREME would reflect artifacts learned by the model. For instance, Enformer has been previously found to underestimate the effects of distal enhancers^6^. In agreement, CREME’s Distance Dependence Test also showed that the effect of enhancer tiles drops off substantially with distance. Notwithstanding, identifying this bias with controlled *in silico* experiments is helpful to debug issues of genomic DNNs, enabling future iterations to address their gaps. Another issue is that the perturbed sequences may introduce an out-of-distribution shift^71^, for which model predictions can be less reliable – not grounded in biology. By staying close to genomic sequences, performing multiple trials, and carefully considering control experiments, we aimed to limit the negative impacts of distributional shifts.

Despite the current limitations of genomic DNNs in comprehensively understanding gene regulation, CREME advances our ability to scrutinize these “black box” models, facilitating the elucidation of the biological factors that drive their predictions. This can also help to understand the strengths and weaknesses of these class of models, informing gaps that can be addressed in future iterations.

### Hyperparameter choices

CREME’s main hyperparameter choice is the tile size and the number of perturbations. In this study, we opted for a tile size of 5 kb as it balances the potential biological insights with the scale of perturbation experiments using a very large Enformer model. Of course, if multiple enhancers and silencer elements fall within a 5 kb tile, then this could affect some of the specific takeaways. Notwithstanding, we demonstrated how a Fine-Tile Search could be used to obtain a higher-resolution view of sequence elements. In addition, the normalization of how the results are presented can skew results to different interpretations. We provided our rationale for the chosen normalizations but alternative choices may be more insightful; this depends on the question being asked. Furthermore, the choice of stratification of the results according to context shuffles was a choice in this study. Further stratification could help to further dissect the rules of gene regulation, especially for highly expressed genes versus lowly expressed genes.

### Hypothesis generation

CREME opens the possibilities to explore what genomic DNNs have learned, moving beyond predictive capabilities to existing experimental datasets and explainable AI methods that make additive approximations of model predictions. Importantly, insights gained through model interpretability should be treated as a hypothesis and not a replacement for laboratory-based experiments. With further advances in machine learning that drive the alignment between genomic DNNs and biological reality to narrow, methods like CREME could facilitate better-informed hypotheses that can help guide more efficient laboratory-based perturbation experiments.

CREME can help prototype experiments and generate plausible hypotheses of *cis*-regulatory mechanisms. By interrogating Enformer, we found that the TSS activity of genes is affected by the complex interactions of multiple enhancers and silencers. This suggests that perturbations with CRISPRi on a single locus (and, in some cases, pairs of loci) would be insufficient to characterize dependencies between CREs fully.

### Moving forward

While this study focuses on the impact of CRE-level perturbations on gene expression predictions within a few cell lines given by Enformer, CREME is general. It can be applied to any sequence-based DNN at any desired resolution. By deploying smaller tile sizes, the experiments performed by CREME can also be used to study motifs and their interactions, even within modestly sized DNNs. CREME provides a roadmap to probe what *cis*-regulatory mechanisms the DNN has learned, moving beyond evaluations based on alignment to experimental data. This toolkit is extensible; different perturbation experiments can be explored to address different biological questions. In the future, we aim to expand CREME to incorporate tests to study enhancer-promoter compatibility rules at both the CRE- and motif-level, study the rules of interactions of weak enhancers and silencers (as opposed to strong enhancers and silencers explored in this study), uncover local *cis*-regulatory networks to identify direct interactions between CREs, and design synthetic CREs that are optimized for a target gene.

## Methods

### Enformer

Enformer is a previously established DNN that takes as input genomic sequences of length 196,608 bp and predicts 5,313 epigentic tracks for human and 1,643 epigenetic tracks for mouse biosamples through two output heads^1^. For each track, Enformer’s predictions cover 896 binned positions, with each bin representing 128 bp. This represents the central 114,688 bp of the input sequence. The extended input sequence, provides context for the edge cases, i.e. the start and end of the predictions. The epigenetic tracks consist of processed coverage values of expression (CAGE), DNA accessibility (DNase-seq), transcription factor binding and histone modification (ChIP-seq). Enformer is composed of convolutional layers that initially summarize the input sequence into representations of 128 bp bins. This is followed by 11 transformer blocks that use multi-head self-attention^72^. We acquired code for the Enformer model along with trained weights from https://tfhub.dev/deepmind/enformer/1 as per instructions in the Methods section of Ref.^1^.

### Borzoi

Borzoi is a DNN similar to Enformer in that it predicts a range of epigenetic tracks from input DNA sequences. However, it considers a larger input size of 524,288 bp and makes predictions for strand specific outputs of CAGE (and other tracks) at a 32 bp resolution. Borzoi is an ensemble of 4 models trained on different data splits. Here, we adopted the same strategy as the authors of Borzoi of running inference using all 4 models and averaging the results. Each model in the ensemble is composed of convolutional layers that summarize the input into representations of 128 bp bins, 8 transformer layers that use multi-head self-attention and deconvolution layers that upsample the representations to 32bp resolution. We acquired code for the Borzoi model along with trained weights from https://github.com/calico/borzoi/tree/main as per instructions in the Methods section of^4^.

### Transcription start site selection - Enformer

We acquired human annotations from GENCODE^25^ (https://www.gencodegenes.org/human/) and filtered for ‘transcript’ annotations and ‘protein coding’ genes. We then extracted sequences of length 196,608 (or 524,288 for Borzoi) from the GRCh38 reference genome centered at each filtered TSS. We converted the sequences to a one-hot representation, treating N characters as a uniform probability (i.e. 0.25). We calculated Enformer’s prediction for these sequences and considered the mean at positions 447 and 448 (of the 896 binned predictions), which corresponds to the central TSS. We used tracks 4,824, 5,110 and 5,111 of the human output head (corresponding to PC-3, GM12878 and K562 CAGE predictions, respectively). We refer to this scalar coverage value per cell line as the TSS activity. To focus our study on genes that yield high TSS activity, we considered the top 10,000 unique genes per cell line with the highest read coverage.

### CREME: *cis*-Regulatory Element Model Explanations

CREME is an *in silico* perturbation assay toolkit that can uncover rules of gene regulation learned by a large-scale DNN. The rationale behind CREME stems from the concept that DNNs are function approximators. Thus, by fitting experimental data, the DNN is effectively approximating the underlying “function of the experimental assay”. By treating the DNN as a surrogate for the experimental assay, CREME can be queried with new sequences and provide *in silico* “measurements”, albeit through the lens of the DNN. Inspired by wetlab experiments, such as CRISPRi^23,24,73^, that perturb genomic loci to uncover how CREs influence gene expression, we devised a suite of *in silico* perturbation experiments that interrogate a DNN’s understanding of long-standing questions of gene regulation, including the context dependence of gene expression^30,74^, identification of enhancing and silencing CREs and their target genes^23,29^, distance dependence of CREs to target genes on gene expression, the complex higher-order interactions of CREs and the effect of finer-scale elements on gene expression^33,34,59–61,65^. Since DNN predictions may not fully capture the underlying biology when fitting experimental data, CREME is strictly a model interpretability tool. Below, we detail the different *in silico* perturbation tests explored in this paper.

### CREME investigation of Enformer

For the vast majority of the experiments, we only considered TSS activity, which we define as the central 5 kb tile centered on the input sequence. Enformer’s receptive field for this tile covers roughly 200 kb sequences, so the 200 kb region centered on the sequence is what is probed in our expeirments. We split the central 200 kb sequences into 38 non-overlapping 5 kb tiles (such that the tiles are fully within the input sequence), with the central tile corresponding to the TSS of an annotated gene. We define the *TSS activity* as the central bin in Enformer’s prediction, i.e. the mean of positions 447 and 448 of tracks 4,824, 5,110 and 5,111 of the human output head.

### Context Dependence Test

The *Context Dependence Test* aims to measure the effect size of TSS activity in random contexts (derived from dinucleotide shuffled versions of the wild type sequence). This test measures the extent to which a prediction of a given TSS activity is influenced by its context which may contain enhancers and silencers. To perform the Context Dependence Test, we executed the following steps:

1. Predict TSS activity for the wild type sequence (denoted as WT).
2. Dinucleotide shuffle the sequence (except the 5kb tile centered at the TSS).
3. Predict TSS activity for the shuffled sequence (denoted as MUTANT).
4. *Normalization:* compute context effect on TSS using WT as control: (WT - MUTANT) / WT
5. Repeat steps 2-4 10 times and average across different random dinucleotide shuffles.

**Example case**. Assuming WT prediction is 10, MUTANT is 5 (i.e., shuffling leads to a drop in activity) the normalized effect would equal:

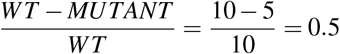

### Interpretation

Effect size of 0 means that the context is neutral and has no effect on the TSS predictions (i.e. wild type and mutant yield the same prediction). Positive effect size means that the central TSS prediction for the mutated sequence is lower than wild type, which indicates that we have perturbed an enhancing context. Negative effect size means that the central TSS prediction for the mutated sequence is higher than wild type, which suggests that we have perturbed a silencing context.

### Analysis

We categorized the sequences into silencing, neutral, and enhancing contexts based on their context effect on TSS. We identified 3 regions: (i) enhancing context were chosen based on an effect size of more than 0.95, (ii) neutral context was chosen if the absolute effect size was less than 0.05, and (iii) silencing context was chosen based on an effect size of less than -0.3. If the number of data points in a category was above 200, we randomly sampled 200 sequences to cap the number for further experiments. We used these groups per cell line throughout the experiments.

For each cell line the breakdown of contexts in each category is given in Supplementary Table 2.

For Borzoi, we chose to proceed with less strict thresholds: - 0.9, 0.05, and -0.2 for enhancing, neutral and silencing contexts, respectively, to address the weaker context effect sizes given by Borzoi. The number of detected sequences in each category is given in Supplementary Table 3.

#### Context Swap Test

Context Swap Test aims to measure the extent that TSS activity depends on a specific genomic context or measure compatibility with other contexts. To perform the Context Swap Test, we executed the following steps:

1. Get the central TSS (5 kb tile) from the source sequence.
2. Insert the source TSS in each of the target sequences at the central TSS position, thereby replacing the existing TSS of the target sequence.
3. Predict TSS activity for the mutant sequence (denoted as MUTANT) and the wild type source sequence (WT).
4. *Normalization:* compute fold change over control according to: MUTANT / WT

**Example case**. Assuming WT prediction for a given sequence from which the TSS is taken is 20, MUTANT, i.e. the activity of the same TSS in a new background is 10 (i.e., moving the TSS to the new background leads to a lower activity) the normalized effect would equal:

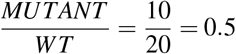

Assuming both sequences are from enhancing backgrounds, this would indicate that the new context is less enhancing compared to its native TSS (classified based on Context Dependence Test).

### Interpretation

If the fold change over control is 1, the TSS activity in the new context is the same as in its native context. If the value is below 1 that means that the TSS is less active in the new context and vice-versa for values above 1.

### Analysis

We performed the Context Swap Test on the sequences filtered by the Context Dependence Test per cell line. Specifically, we placed the TSSs in each context category across all other context categories, separately keeping track of the source TSS and the context category.

#### Necessity Test

The Necessity Test measures the importance of a putative CRE on the central TSS activity for a given sequence context while the other tiles remain intact. To perform the Necessity Test, we executed the following steps:

1. Predict TSS activity for the wild type sequence (WT).
2. For each 5 kb tile not overlapping with the central TSS:
  a. Dinucleotide shuffle the 5 kb tile under investigation.
  b. Predict TSS activity for the shuffled sequence (MUTANT).
  c. Repeat 10 times and calculate the mean TSS activity.
3. *Normalization:* compute the normalized shuffle effect as: (WT - MUTANT) / WT

**Example**. Assuming WT activity is 10 and it drops to 5 after shuffling 1 CRE (indicating that the shuffled CRE is enhancing), the normalized shuffle effect would equal:

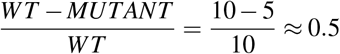

### Interpretation

Effect size of 0 means that the WT value is similar to the case when the tile is shuffled, and therefore, the tile shuffle has no effect on the TSS. A large positive value means that the shuffled case yields a lower TSS signal than the WT, i.e. the tile shuffling leads to a drop in TSS activity and a large negative value means the shuffling leads to a higher TSS activity compared to WT.

### Analysis

We performed the Necessity Test on all tiles within the sequences from Context Dependence Test that had enhancing, silencing or neutral backgrounds (as classified by selected thresholds).

### Sufficiency Test

The Sufficiency Test measures the effect of a given CRE on its TSS in otherwise random contexts, i.e. in isolation from the rest of the tiles from the original wild type sequence. This essentially measures whether the CRE by itself is enough to up or downregulate the TSS. To perform the Sufficiency Test, we executed the following steps:

1. Predict TSS activity for the wild type sequence (WT).
2. Dinucleotide shuffle the sequence.
3. Add the TSS 5 kb tile and predict TSS activity (CONTROL).
4. Add the CRE and the TSS tiles to the sequence and predict TSS activity (MUTANT).
5. *Normalization:* We used 2 different normalizations - one for enhancing context sequences and one for the neutral or silencing context sequences. This is motivated by the different activity levels of TSS we get in the CONTROL case (enhancing context TSSs by definition have very low activity when the context is shuffled). (i) in case of enhancing context sequences, we compute the normalized CRE effect as (MUTANT - CONTROL) / WT. (ii) in case of neutral and silencing context sequences, we compute (MUTANT - CONTROL) / CONTROL.
6. Repeat each shuffle 10 times and average the normalized CRE effect per sequence.

**Example**. Assume an enhancing sequence with a WT prediction of 10, CONTROL (only TSS) sequence with predicted activity of 2. Given a strong enhancer CRE that recovers the activity in the MUTANT (TSS and CRE) to 12, the normalized activity would equal:

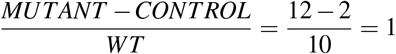

Given a silencing context sequence with WT activity of 10, CONTROL activity of 20 (note that silencing context sequences by definition have higher activity after context shuffle) and a strong silencer that represses the signal to 10 the normalized effect would be computed as:

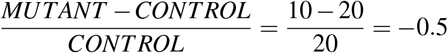

### Interpretation

In case of enhancing context sequences, the TSS activity drops to a small value after the whole context shuffling (CONTROL). Therefore, here we compare the value to the original WT sequence, subtracting the small activity of the TSS on its own (CONTROL). This can be interpreted as the extent to which a given tile individually restores TSS activity to the original WT value. A value of 0 means that the tile has no positive effect on the TSS (MUTANT equals CONTROL), positive values indicate an enhancing effect (MUTANT > CONTROL) and vica versa for negative values.

In case of silencing and neutral context sequences, the TSS activity in CONTROL condition is not always a small value (by definition the silencing contexts are the ones where shuffling the context leads to a high TSS activity). This leads to ambiguities in cases when the WT and CONTROL contribute to the normalized values. Therefore, we simply computed the tile effect as fraction change observed when adding the tile compared to CONTROL. Value of 0 means that tile addition has no effect on TSS, a positive value indicates that tile addition activates the TSS and a negative value means that the tile lowers TSS activity.

#### Analysis

We performed the Sufficiency Test on the same subset of sequences as Necessity Test that had enhancing, silencing or neutral backgrounds (as classified by selected thresholds). Based on Sufficiency Test results, we denote CREs from enhancing contexts with effect size larger than 0.3 as enhancers. Similarly, we define silencers as tiles from silencing contexts with effect size smaller than -0.3. In case of Borzoi, we used -0.15, a higher threshold, for silencers to include more elements and considered those in both silencing and neutral background.

#### TSS-CRE Distance Test

TSS-CRE Distance Test is a GIA experiment where we systematically shift the position of a tile in shuffled sequences and measure its effect on TSS activity. For each putative CRE tile, we performed the TSS-CRE Distance Test by executing the following steps:

1. Dinucleotide shuffle the sequence and embed the central TSS 5 kb tile.
2. Define test positions P as non-overlapping 5 kb tile start positions such that each tile is fully included in the window of the model input (excluding the TSS tile).
3. For each test position P:
  a. Insert the CRE tile at position P in the dinucleotide shuffled sequence (with an intact TSS) and predict TSS activity (MUTANT).
  b. Define CONTROL as the sequence with maximum TSS activity among the MUTANT sequences.
  c. *Normalization:* Compute the fold change over control as MUTANT / CONTROL
  d. Repeat each shuffle 10 times and average the fold change over control per sequence.

**Example**. Assuming we consider only 5 possible test positions with MUTANT sequence activity values of 2, 5, 10, 5, 2 the CONTROL would be set to 10 and the normalized activities would equal 0.2, 0.5, 1, 0.5 and 0.2 respectively.

#### Analysis

We used the definition of enhancers and silencers based on Sufficiency Test results. We performed the TSS-CRE Distance Test on CREs defined as enhancers within enhancing contexts and silencers in silencing contexts for each cell line.

#### Higher-order Tile Interaction Test

The aim of Higher-order Tile Interaction Test is to dissect CRE networks. Specifically, we compute the combined effect of multiple tile shuffles that have large effects through a greedy search. For enhancers, the iterative greedy search systematically identifies tiles that lead to a lower TSS activity when shuffled. We followed the same steps for silencer search but instead of choosing the minimum predicted value we chose the maximum predicted value in each iteration. To perform the Higher-order Tile Interaction Test, we executed the following steps:

1. Predict TSS activity for the wild type sequence (WT).
2. For each greedy search iteration:
  a. For each tile that is not fixed (i.e. fixed tiles are the central TSS tile and tiles selected from previous rounds):
    i. Dinucleotide shuffle the tile
    ii. Predict TSS activity for the mutant sequence (MUTANT)
    iii. *Normalization:* Compute the fold change over wild type, i.e. MUTANT/WT
    iv. Repeat each shuffle 10 times and average normalized output per sequence.
  b. Fix the tile that yields the maximum effect on TSS activity. For enhancers, maximal decrease in TSS activity; for silencers, maximal increase in TSS activity. The shuffled version that is most representative is chosen when fixing. This is selected based on the instance that yields a prediction closest to the mean across 10 shuffles.
  c. Repeat for the desired number of rounds in the greedy search or until the entire sequence is fixed.

**Example**. Assume a sequence with 2 enhancing tiles E1 and E2 and WT activity 100. In the first iteration after shuffling E1 the prediction drops to 50, shuffling E2 yields a prediction of 70 while shuffling the other tiles maintains activity at WT levels. The normalized TSS activity in case of shuffling E1 would be computed as:

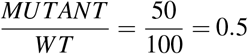

Given that this is the sharpest drop in TSS activity, if searching for enhancers this tile would be fixed or shuffled.

#### Analysis

We performed Higher-order Tile Interaction Test for maximally enhancing TSS activity and maximally silencing TSS activity for all sequences from different context categories.

### Comparison to additive effects

To help understand the trajectories from the Higher-order Tile Interaction Test, we calculated the hypothetical effects of an additive model. In brief, the additive effects are calculated based on combining the effects on TSS activity from the individual effects of each CRE (i.e. calculated in the first round of the greedy search), following the CRE tile order found by the greedy search. This does not take into account cooperative or redundant relationships within sets of CREs, which would be captured in the greedy search.

To compute the hypothetical additive effects, we performed the following steps:

1. Predict TSS activity for the wild type sequence (WT).
2. Denote the order of tile fixing or shuffling from greedy search results of iteration 1 as T.
3. Compute effect sizes of individual tile shuffling as MUTANT - WT.
4. Sort the effect sizes according to the tile order T.
5. Following the tile order T, calculate the cumulative sum of the individual tile effects (M_additive). This assumes an additive model.
6. *Normalization:* compute the hypothetical fold change over control according to (M_additive)/WT.

**Example**. Assuming a sequence with 2 strong enhancers E1 and E2 with fully additive effects we would expect their effect sizes to add up to the prediction from when both are shuffled simultaneously. For instance, if WT prediction is 100, and in iteration 1 shuffling just E1 leads to prediction of 60, E2 of 70 then the effect sizes are -40 and -30 respectively. The hypothetical additive model would then predict that the simultaneous shuffling of E1 and E2 would lead to a normalized effect size of:

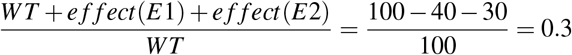

### Interpretation

If the hypothesis of additive effect holds, we would expect the tile greedy search trace to be the same as the additive or hypothetical trace for each sequence. If the results are different we can categorize these as superadditive or subadditive for cases when the additive model overestimates the shuffle effect or underestimates it, respectively. To illustrate an example scenario leading to a superadditive case let us assume two enhancers are cooperating, i.e. their combined effect is larger than individual effects (a non-additive case). We would expect their individual shuffle effects to also be larger than shuffling them simultaneously (because disabling one leads to a large effect size already). In contrast, subadditivity can arise if two enhancers are redundant, i.e. their roles are overlapping, the effect size will be small when only a single tile is shuffled (because the other enhancer tile can compensate). Therefore, the estimated additive effect (based on single tile shuffles of iteration 1) will underestimate the effect of shuffling both of the enhancers. This will thus lead to the additive hypothetical trace being higher than the one based on the greedy search.

#### Analysis

To characterize deviations, we computed the mean squared error (MSE) of the greedy and hypothetical additive outputs for each sequence. We classified the cases where the MSE value is above 0.1 (arbitrary threshold) and the greedy search results on average is greater than the average of additive as superadditivity cases. Similarly, we classified the cases where the MSE value is above 0.1 (arbitrary threshold) and the greedy search result on average is lower than the average of additive as subadditivity cases. We used a stricter threshold of 0.05 for the MSE value to classify sequences as additive. We applied these threshold to both the enhancer and silencer search cases after assessing visually that the traces overlap substantially.

### Comparison to multiplicative effects

Similarly, we compared greedy search outcomes to a multiplicative model. For this we computed the natural logarithm of all the values used and otherwise proceeded with the same steps as in *Comparison to additive effects*. We interpreted the results using the same logic as in the additive effect case, but we used different thresholds for classifying sequences into multiplicative and non-multiplicative categories. We mostly observed cases of the hypothetical model predicting higher values than the greedy search yielded and some where they aligned (hence the binary classification). We used the MSE thresholds 0.04 for enhancers (cases of MSE above 0.04 were classified as non-multiplicative) and 0.01 for silencers. Both were selected after visual inspection of the traces.

### Sufficiency of CREs identified through higher-order interaction test

In a complementary set of experiments to higher-order interaction test, we tested if the identified CREs are sufficient to recover or suppress the TSS activity. To do this we performed the following steps:

1. For a given sequence retrieve the order of CRE shuffling from the higher-order interaction test.
2. Dinucleotide shuffle the sequence and embed the central TSS tile (CONTROL).
3. Sequentially embed consecutive enhancer (or silencer) CREs identified during the higher-order interaction test (MUTANT).
4. In each step compute the normalized TSS activity - for enhancing context sequences as MUTANT / WT, for silencing context sequences as MUTANT / CONTROL.

### Multiplicity Test

The Multiplicity Test measures how TSS activity scales upon repeated addition of an enhancing or silencing tile. With this GIA experiment, we aim to test the model’s extrapolation behavior. Specifically, we probed whether TSS activity reaches saturation upon a high dosage of a CRE; saturation is when the predictions reach a plateau when we enrich for enhancers or silencers. The Multiplicity Test is similar to the greedy search used in the Higher-order Tile Interaction Test, with the exception that we are systematically adding the same CRE of interest into optimal positions in each round of dinucleotide shuffled sequences. To compute the Multiplicity Test, we performed the following steps:

1. Define the CRE tile and number of times the CRE will be inserted (number of iterations).
2. Dinucleotide shuffle the sequence while maintaining the central TSS tile intact.
3. Add the CRE of interest in its original position and predict TSS activity (CONTROL).
4. Define test positions as non-overlapping 5 kb tile start positions such that each tile is fully included in the window of the model input (excluding the TSS tile).
5. Systematically measure predicted TSS activity when CRE is embedded at each unfixed position (MUTANT_POSITION).
6. *Normalization:* calculate the fold change over control given by MUTANT_POSITION / CONTROL
7. Fix the CRE at the position where it maximally affects TSS activity.
8. Repeat steps 3 to 6 for the defined number of iterations.

**Example**. Assuming CONTROL activity of 10, if inserting CRE at a new position yields prediction of 20, the normalized TSS activity would be computed as:

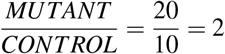

#### Analysis

Using the enhancers defined in Sufficiency Test, we performed 15 iterations of the Multiplicity test for each CRE.

### Biochemical characterization of CREs

To characterize and contrast the chromatin state of the putative enhancing and silencing CREs (at 5 kb window size) we downloaded ENCODE bigwig files and summarized the values at the coordinates of the CREs (Supplementary Data 1). To do this we followed these steps for each cell line:

1. Filter the ENCODE metadata table for given cell line, genome assembly GRCh38, file status of ‘released’ and normalized by ‘fold change over control’ (for tracks other than DNase-seq).
2. Filter, download and classify tracks into (i) histone marks (ii) accessibility (i.e. DNase-seq, ATAC-seq) (iii) TF binding.
3. Compute the average values if more than one bigwig is available for the same assay and target (e.g. CTCF ChIP-seq) in the same cell line.
4. For each histone and accessibility track, we computed the mean bigwig values across the genomic ranges for the corresponding putative CRE tiles. For TFs, we computed the maximum signal of the bigwig values.

We computed the percentage of enhancer and silencer sequences containing repeating sequences using RepeatMasker using the default parameters. The detailed outputs per repeating sequence category and cell line are given in the Supplementary Table 1.

### Fine-Tile Search

This analysis aims to identify subsets of sequences that recover the majority of a given 5 kb enhancer CRE effect. To formalize the problem, we aimed to identify the smallest set of 50 bp sequences that, when embedded in otherwise shuffled context, recovers at least 70% of the enhancer CRE effect on the TSS. We did this using a two-stage nested greedy search, reducing the search space in each stage.

1. Dinucleotide shuffle the sequence, embed the TSS and enhancing 5 kb CRE (at the original position), compute mean prediction and denote this as CONTROL (repeat 10 times).
2. Perform greedy search using non-overlapping window size of 500 bp; dinucleotide shuffle 500 bp windows (one at a time) and select the tile that leads to minimal reduction in TSS activity (MUTANT). Compute CRE effect recovered ratio as MUTANT / CONTROL and repeat while this fraction is above 0.9.
3. Perform greedy search using non-overlapping window size of 50 bp; search and shuffle set of 10 fine-tiles of size 50 bp so that the TSS activity drops minimally (MUTANT). Compute CRE effect recovered ratio as MUTANT / CONTROL and repeat while this fraction is above 0.7.

We compared this approach to XSTREME^51^ (using both *de novo* motifs and known motifs from the JASPAR database) and saliency-based motif embedding. For XSTREME analysis, we scanned the wild type sequence for motif hits with FIMO^75^ and embedded those subsequences into otherwise shuffled background sequences. We evaluated how much of the CRE effect is recovered by XSTREME motifs using the same normalized value (i.e., by dividing with the activity when the whole CRE is embedded). For saliency-based fine-tile analysis, we did the following:

1. Compute saliency (gradient*input) for the 2 central output bins for the given cell line track.
2. Sort the values in descending order.
3. Use the same CONTROL sequences as in CREME fine-tile search (to make the results directly comparable).
4. For values of N ranging from 500 to 5000 in increments of 500bp embed the first N base-pairs into shuffled background sequences (MUTANT).
5. Compute CRE effect recovered ratio as MUTANT/CONTROL.

#### Analysis

For this analysis we used all the enhancing context sequences and the enhancer CREs identified within those in each cell line (74 in K562, 41 in GM12878 and 35 in PC-3. We used the same set of 10 backgrounds per sequences for comparing CREME, XSTREME and saliency-based analyses.

## Data Availability

Results from this paper is deposited at zenodo: doi.org/10.5281/zenodo.8111754.

## Code Availability

Open-source code to deploy CREME can be found at GitHub: https://github.com/p-koo/creme-nn. The code for reproducing the analyses in the manuscript is available at GitHub: https://github.com/shtoneyan/CREME.

## Acknowledgements

The authors would like to thank Saket Navlakha, Jack Desmarais, and members of the Koo Lab for helpful comments on the manuscript. Research reported in this publication was supported in part by the National Human Genome Research Institute of the National Institutes of Health under Award Number R01HG012131, the National Institute Of General Medical Sciences of the National Institutes of Health under Award Number R01GM149921, and the Simons Center for Quantitative Biology at Cold Spring Harbor Laboratory. This work was performed with assistance from the US National Institutes of Health Grant S10OD028632-01. We would also like to thank the support of the NVIDIA GPU Grant Program.

## Author contributions

ST and PKK conceived of the method and designed the experiments. ST developed code, ran the experiments, and analyzed the results. ST and PKK interpreted the results and contributed to writing the paper.

