## Extended data for "Interpreting *Cis*-Regulatory Interactions from Large-Scale Deep Neural Networks for Genomics"

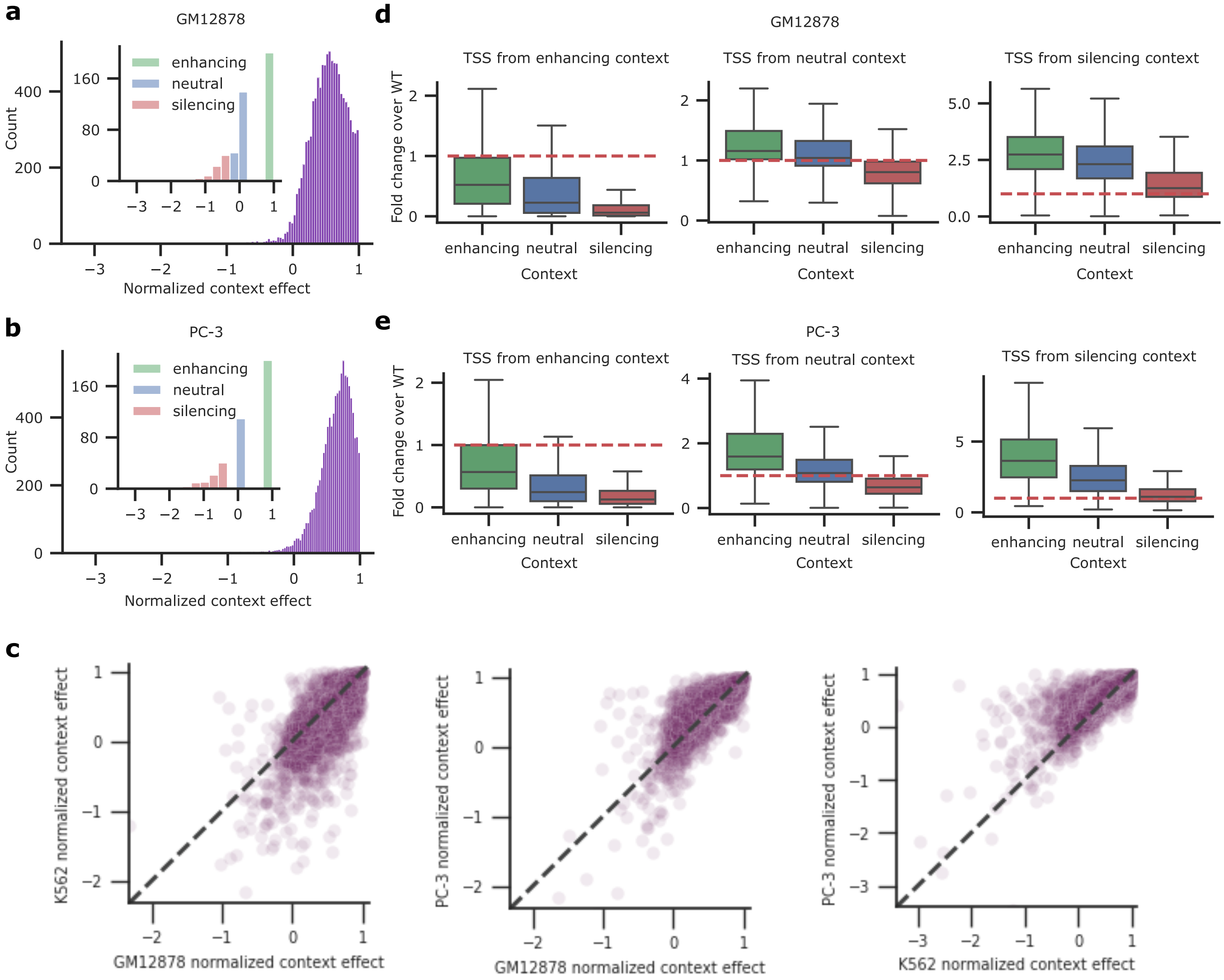


**Extended Data Figure 1.** Results of the Context Dependence Test and Context Swap Test for GM12878 and PC-3. **a, b**, Histogram of normalized context effect from the Context Dependence Test for 10,000 sequences that contain an active, annotated gene in GM12878 and PC-3 cells. Inset shows the subset of sequences for enhancing, silencing and neutral contexts. **a** inset contains 200, 78 and 183 data points in enhancing, silencing and neutral context respectively. **b** inset contains 200, 90 and 110 data points in enhancing, silencing and neutral context respectively. **c**, Pairwise comparison of normalized context effects between cell lines for matched genes. The number of data points is 7688, 6946, 7492 from left to right. **d, e**, Context Swap Test results. Boxplots of normalized context effect on TSS for sequences with context perturbations given by insertion of the original TSS in different context categories. Results are organized according to the original TSS category: enhancing (left), neutral (middle), and silencing (right). The number of data points in each boxplot represent an all-vs-all comparison of each respective TSS in each possible context. The number of data points in **d** is 40,000, 36,600, 15,600 in boxplots for TSS from enhancing context, 36,600, 33,489, 14,274 in TSS from neutral context and 15600, 14274, 6084 in TSS from silencing context. The number of data points in **e** is 40,000, 22,000, 18,000 in boxplots for TSS from enhancing, 22,000, 12,100, 9,900 in TSS from neutral context and 18,000, 9,900, 8,100 in TSS from silencing context. Boxplots show the first and third quartiles, the median (central line) and the range of data with outliers removed (whiskers).

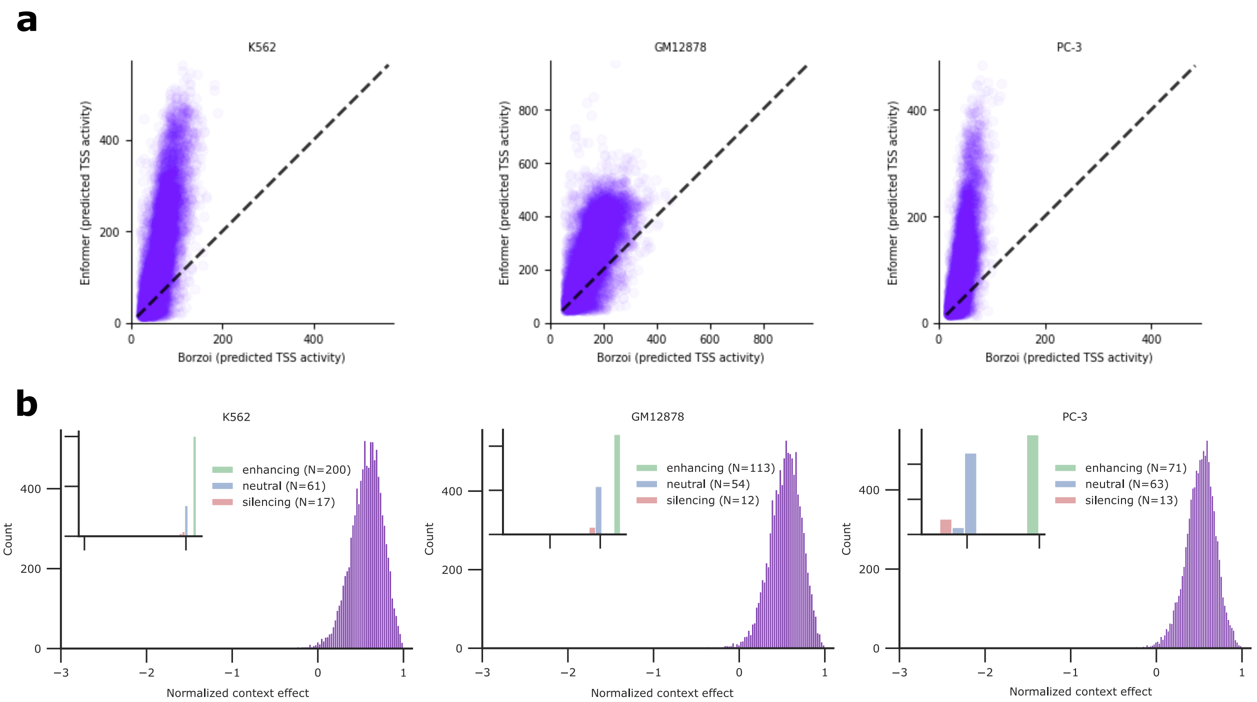


**Extended Data Figure 2.** Borzoi Context Dependence Test results. **a**, Scatter plot comparing the wild-type activity predicted by Enformer versus Borzoi for the matched cell types and for matched genes. **b**, Histogram of normalized context effect for the 10,000 highest activity, annotated genes (according to Borzoi's predictions) for K562, GM12878 and PC-3 cells. Inset shows the subset of sequences for enhancing, silencing and neutral contexts. The number of data points is shown in inset legend.

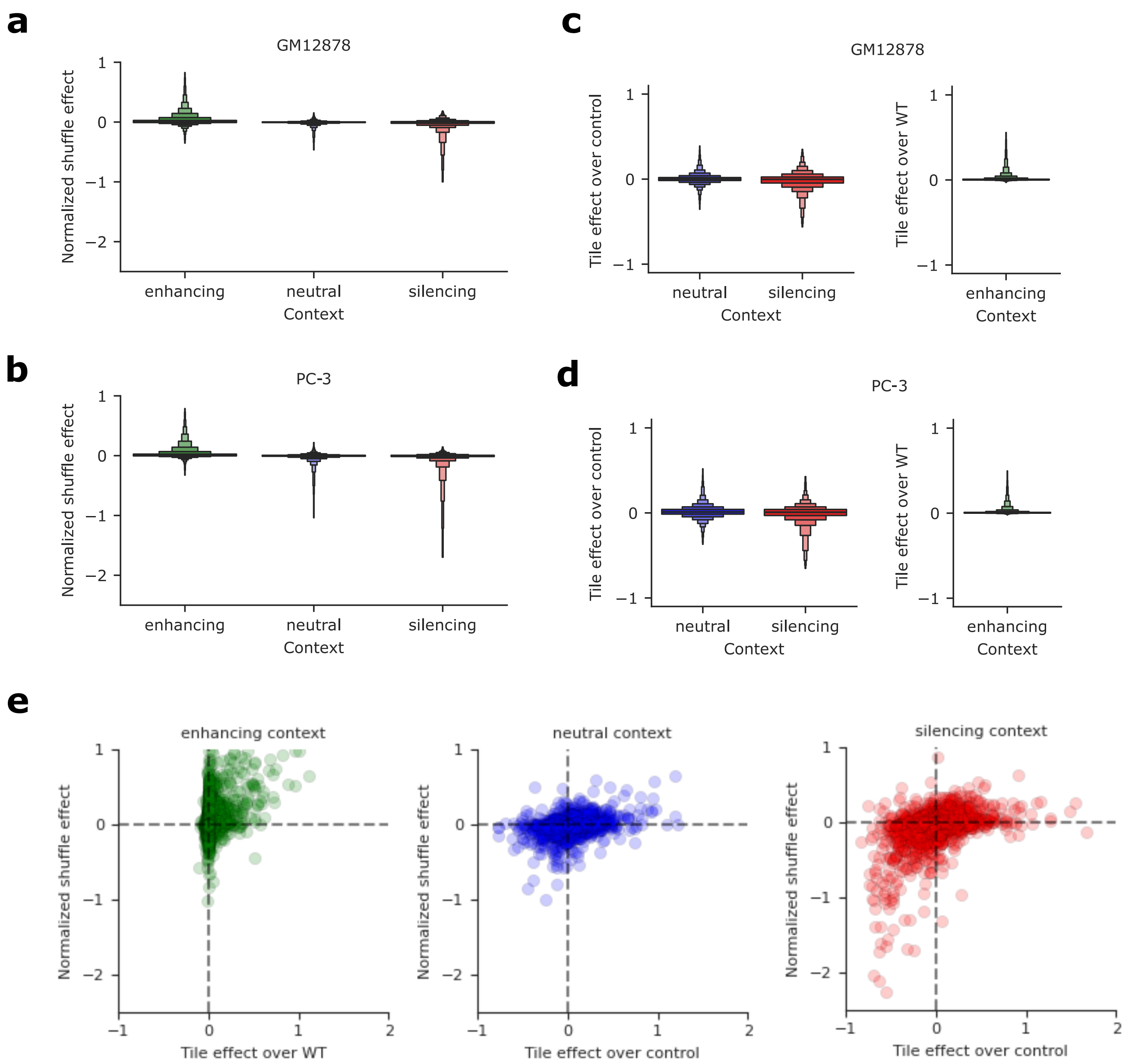


**Extended Data Figure 3.** CRE effects on TSS activity in GM12878 and PC-3 cell lines. **a,b**, Boxen plot of the normalized shuffle effect for each tile in sequences from enhancing, neutral and silencing context categories (Necessity Test) for GM12878 (**a**) and PC-3 (**b**). The number of data points in **a** is 7600, 6954, 2964 and in **b** is 7600, 4180, 3420 in enhancing, neutral and silencing contexts respectively. **c, d**, Boxen plot of tile effects for each tile in sequences from enhancing, neutral and silencing context categories (Sufficiency Test) for GM12878 (**c**) and PC-3 (**d**). Normalization is with predicted TSS activity for wild-type (enhancing context) and control, i.e. the intrinsic TSS activity (neutral and silencing context). Boxen-plots have the same number of data points as in **a** and **b**. **e**, Scatter plot between the results from the Necessity Test (*y*-axis) versus the results from the Sufficiency Test (*x*-axis) in K562 cell line (N = 7,600 in each plot corresponding to 200 sequences with 38 tiles in each).

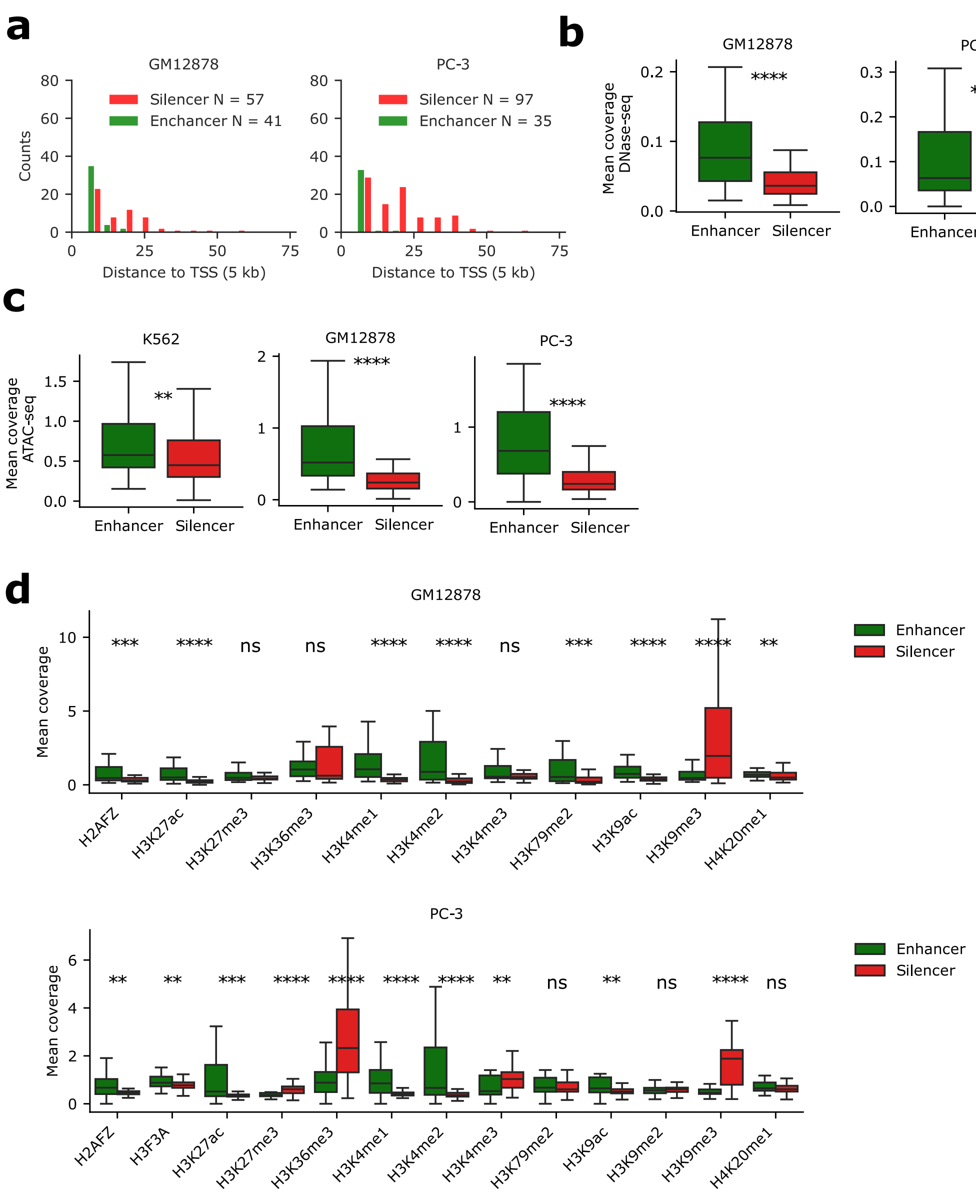


**Extended Data Figure 4.** Characterization of sufficient CREs in GM12878 and PC-3. **a**, Histogram of the distance between CRE tiles from TSS for sufficient enhancers and silencers in GM12878 and PC-3. **b-d**, Boxplots of mean DNase-seq coverage (**b**), mean ATAC-seq coverage (**c**), and mean histone mark coverage (**f**) of sufficient enhancer and silencer tiles in various cell types. The number of points in green and red boxes is 41 and 57 for GM12878 and 35 and 97 for PC-3. Significance is given by the Mann–Whitney U test (*: $p < 0.05$; **: $p < 0.05$; ***: $p < 0.001$; ****: $p < 0.0001$).

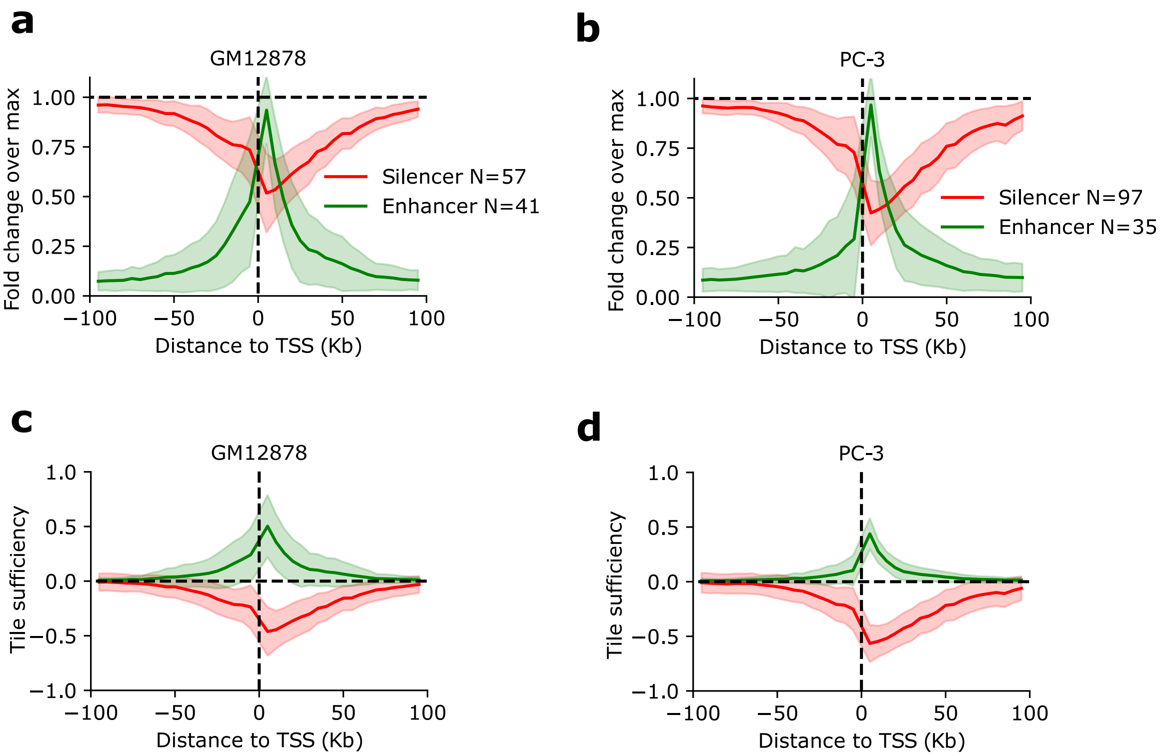


**Extended Data Figure 5.** TSS-CRE Distance Test results across cell lines. **a**-**c**, Average plot of the fold change over max versus distance to TSS for GM12878 (**a**) and PC-3 (**b**). Max represents the maximum TSS activity across all embedded positions within each sequence using Enformer. Shaded region represents standard deviation of the mean. **c**,**d**, Plot of the tile sufficiency versus distance to TSS for GM12878 (**c**) and PC-3 (**d**), respectively. Tile sufficiency is calculated according to the predicted TSS activity with a TSS-CRE pair at a given distance minus the control sequence (shuffled context with just the TSS) divided by the WT sequence for enhancers and by the control sequence for silencers.

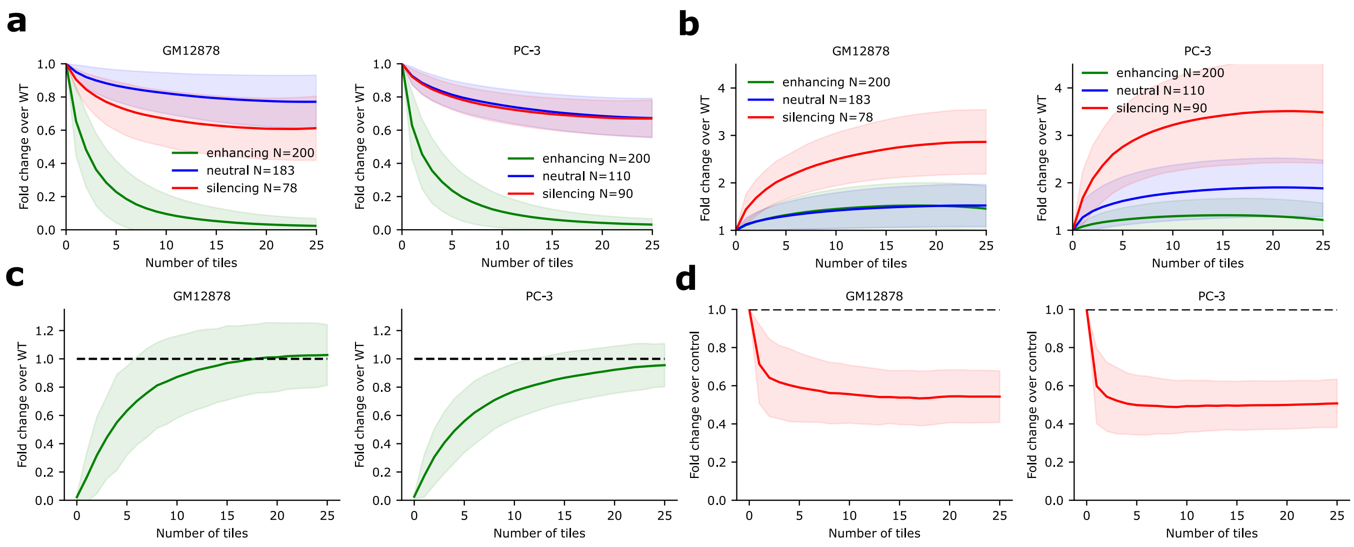


**Extended Data Figure 6.** Optimal CRE sets reveal complex interactions in GM12878 and PC-3. **a**,**b**, Average plot of the greedy search results for enhancer tile sets (**a**) and silencer tile sets (**b**) for sequences from different context categories for various cell lines. The fold change over wild-type (WT) is the predicted TSS activity of the shuffled CRE tiles in each round of the greedy search (indicated by the number of tiles). **c**,**d**, Sufficiency of the tile sets identified in each round of greedy search. Average fold change over wild-type (**c**) and control (**d**), which represents shuffled sequences with just the TSS tile. Sufficiency places the tile sets along with the TSS tile into shuffled sequences, averaging over 10 total shuffles. Shaded region represents the standard deviation of the mean.

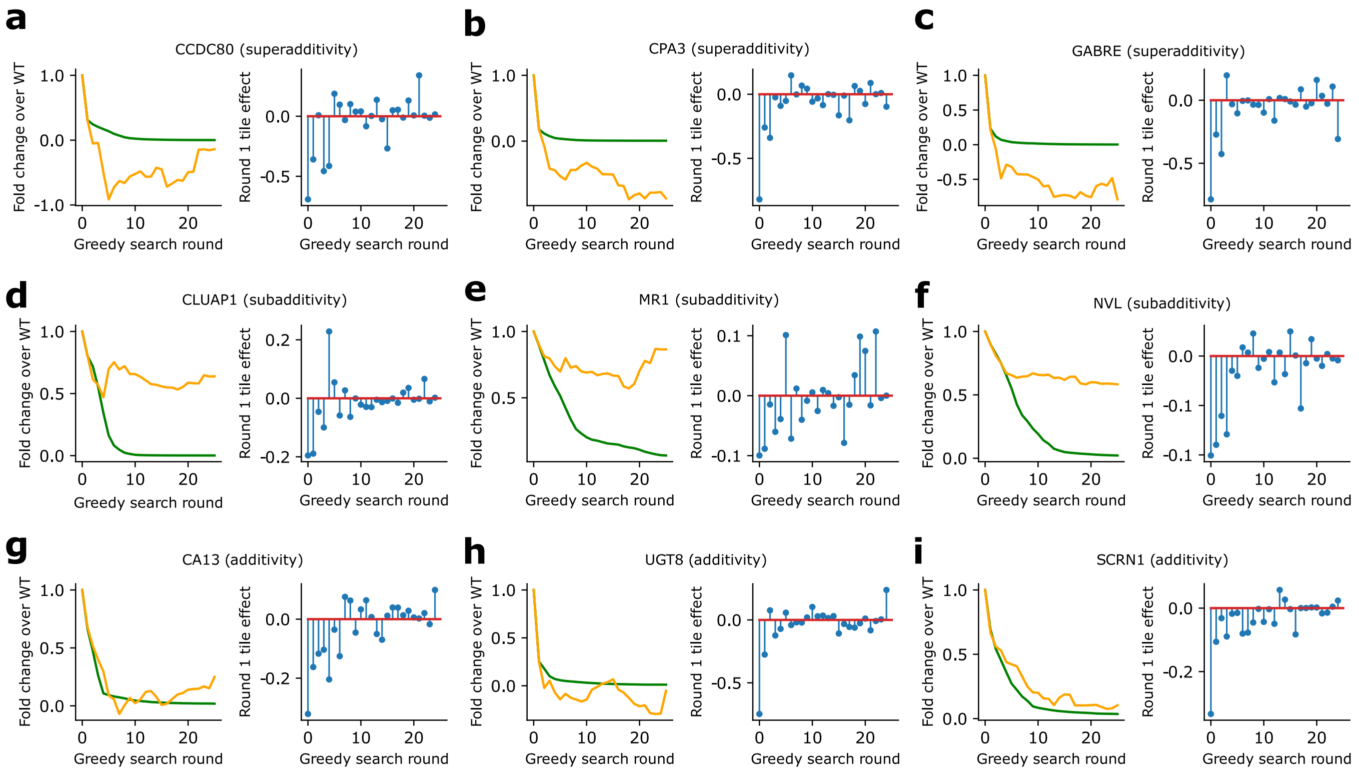


**Extended Data Figure 7.** Example sequences showing individual tile effect sizes from the Higher-Order Interaction Test results. **a**-**i**, the left panels show results of the greedy search (green) and the additive model (orange) for a particular gene; the right panel shows the independent tile effect size (calculated from the first iteration) sorted according to greedy search tile order. **a**-**c** shows example sequences classified as superadditivity; **d**-**f** shows sequences classified as subadditivity; **g**-**i** shows example sequences classified as additivity.

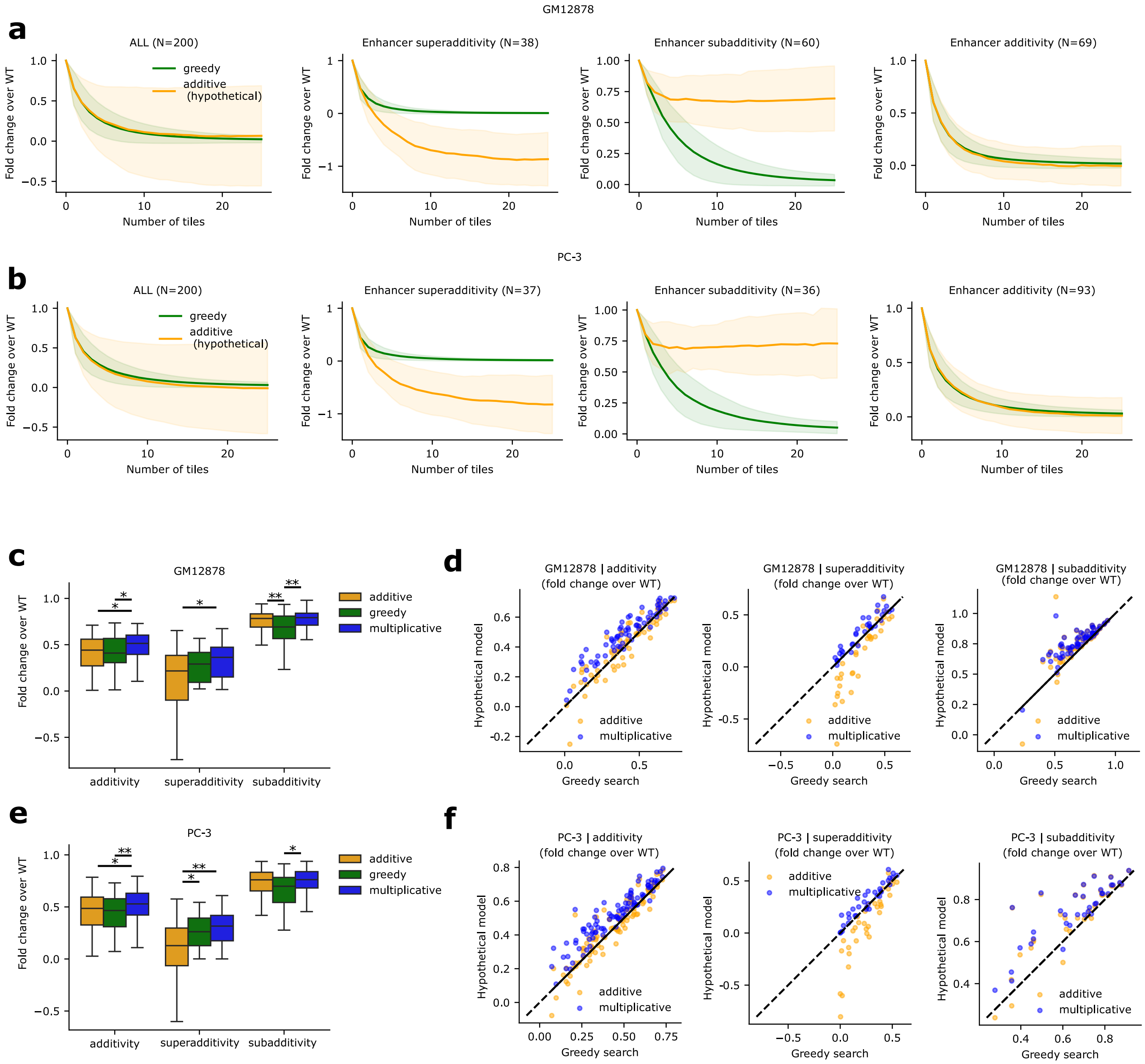


**Extended Data Figure 8.** Comparison of enhancer sets identified by the Higher-Order Interaction Test and a hypothetical additive model for GM12878 and PC-3. **a**,**b**, Comparison of the average fold change over wild-type (WT) for enhancer sets for sequences categorized as enhancing context versus a hypothetical additive effects model. The sequences from enhancing contexts are stratified according to interaction type, superadditivity, subadditivity, and additivity. Sequences were classified using mean squared error based thresholds of 0.1 for superadditivity and subadditivity and 0.05 for additivity definition (with some ambiguous cases left out of classification). Shaded region represents standard deviation of the mean. **c**,**e**, Comparison of hypothetical additive model and hypothetical multiplicative model versus greedy search outcomes at iteration 2 of the higher-order interaction test. The number of points in each box is 69, 38 and 60 in GM12878 and 93, 37, 36 in PC-3 for additive, superadditivity and subadditivity cases. Note, that some ambiguous cases were left out of the classification if they were outside of the selected thresholds. Statistical significance was given according to the Mann-Whitney U test (*: $p < 0.05$; **: $p < 0.01$; ***: $p < 0.001$; ****: $p < 0.0001$). greedy search versus hypothetical additive or multiplicative models. Scatter plots show a more detailed view of the data in **c**,**e** with x-axis showing the higher-order interaction test outcomes and the y-axis showing the hypothetical model outputs (additive or multiplicative).

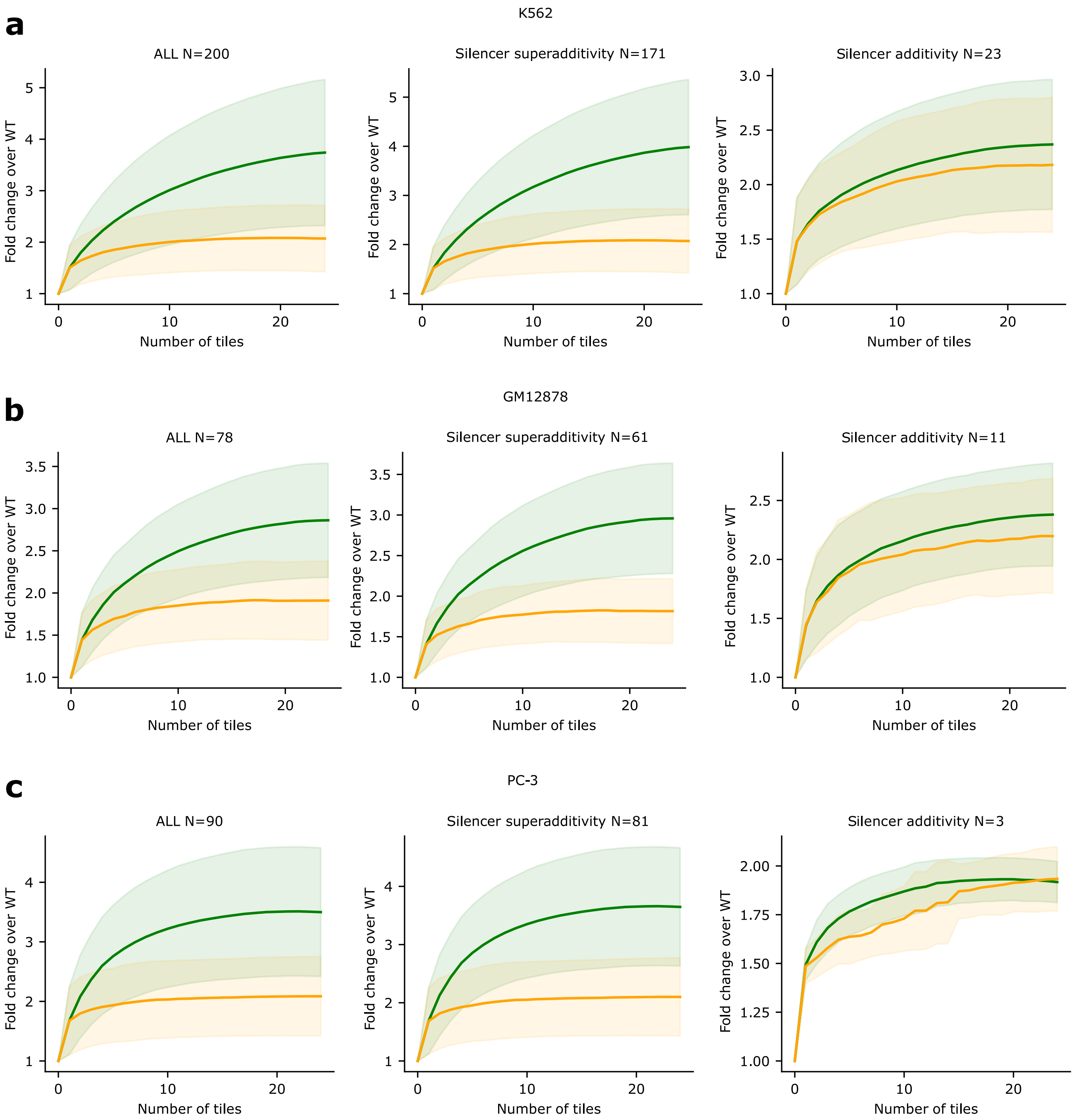


**Extended Data Figure 9.** Comparison of silencer sets identified by the Higher-Order Interaction Test and a hypothetical additive model for K562, GM12878 and PC-3. **a-c**, Comparison of the average fold change over wild-type (WT) for silencer sets for sequences categorized as silencing context versus a hypothetical additive effects model for K562 (**a**), GM12878 (**b**), PC-3 (**c**). The sequences from silencing contexts are stratified according to interaction type, superadditivity and additivity. Shaded region represents standard deviation of the mean. Notably, we did not identify any subadditivity cases.

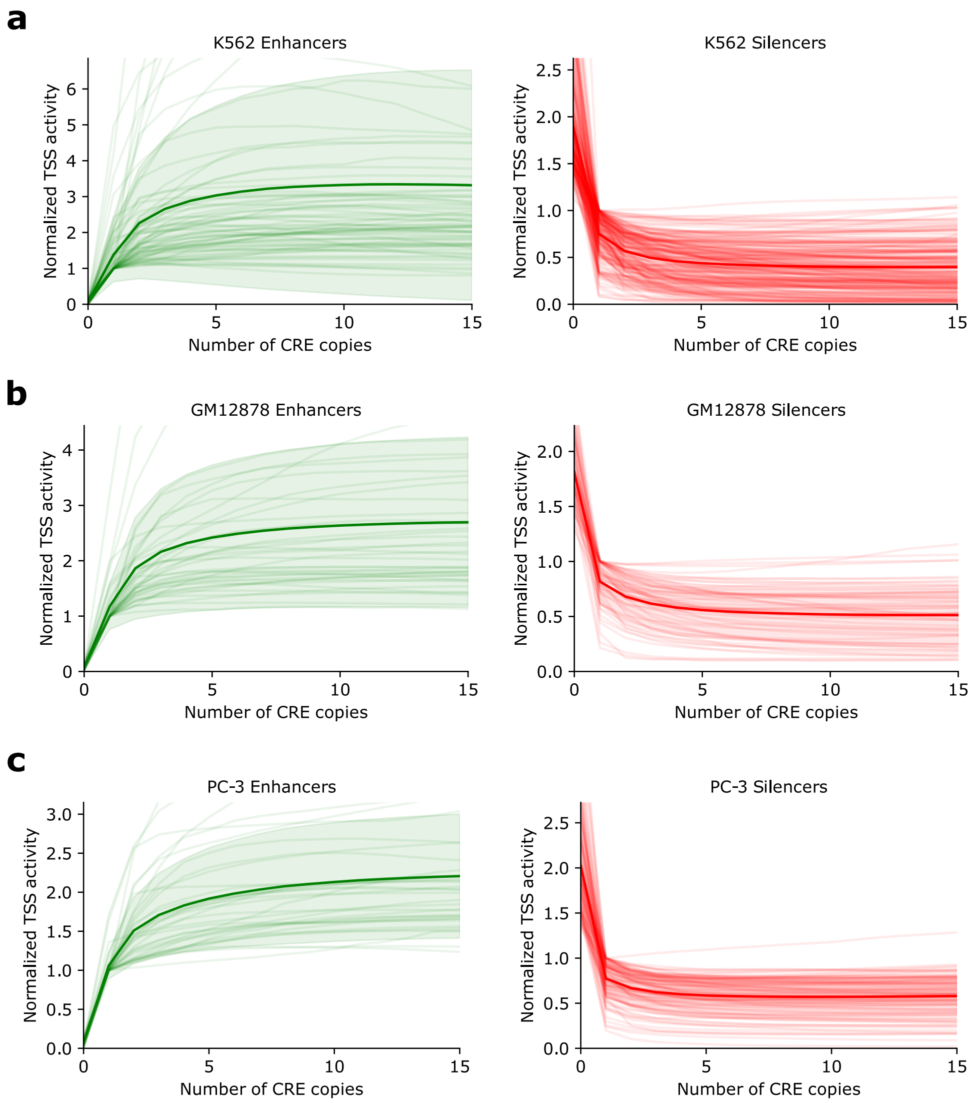


**Extended Data Figure 10.** Saturation behavior of TSS activity predictions by Enformer in various cell lines. The results from a CRE Multiplicity Test applied to sequences from enhancing context (left) and silencing context (right) in **a**-**c**. Each line represents a particular enhancer or silencer CRE embedded into shuffled sequences at optimal positions (according to a Greedy Search) versus the copy number of the CRE in the sequence. The number of enhancers in each plot in **a** - **c** is 200, the number of silencers is 200, 78, 90 in **a** - **c**, respectively. The normalized TSS effect represents the predicted TSS activity of the mutated sequence divided by the control, which is the shuffled sequence with the TSS tile and the CRE in their original positions. The average across all CREs is shown with a thicker line and the shaded region represents the standard deviation of the mean.
