## Supplementary Figures for "Interpreting *Cis*-Regulatory Interactions from Large-Scale Deep Neural Networks for Genomics"

| | K562 | | GM12878 | | PC-3 | |
|---|---|---|---|---|---|---|
| | Enhancer (%) | Silencer (%) | Enhancer (%) | Silencer (%) | Enhancer (%) | Silencer (%) |
| SINEs | 19.66 | 17.08 | 20.34 | 10.69 | 17.33 | 12.22 |
| LINEs | 9.73 | 16.57 | 6.28 | 15.64 | 5.76 | 11.60 |
| LTR | 2.64 | 7.37 | 2.04 | 4.49 | 1.45 | 6.28 |
| DNA elements | 3.24 | 3.15 | 1.75 | 3.89 | 3.27 | 2.64 |

**Supplementary Table 1.** Average percentage that Enformer-defined enhancer or silencer tiles correspond to a repeat element for different cell lines.

| cell line | context category | Number of contexts | Number of contexts after sampling |
|---|---|---|---|
| GM12878 | enhancing | 409 | 200 |
| GM12878 | neutral | 183 | 183 |
| GM12878 | silencing | 78 | 78 |
| K562 | enhancing | 482 | 200 |
| K562 | neutral | 295 | 200 |
| K562 | silencing | 318 | 200 |
| PC-3 | enhancing | 524 | 200 |
| PC-3 | neutral | 110 | 110 |
| PC-3 | silencing | 90 | 90 |

**Supplementary Table 2.** Number of sequences selected in each context category per cell line using Enformer.

| cell line | context category | Number of contexts |
|---|---|---|
| GM12878 | enhancing | 113 |
| GM12878 | neutral | 54 |
| GM12878 | silencing | 12 |
| K562 | enhancing | 204 |
| K562 | neutral | 61 |
| K562 | silencing | 17 |
| PC-3 | enhancing | 71 |
| PC-3 | neutral | 63 |
| PC-3 | silencing | 13 |

**Supplementary Table 3.** Number of sequences selected in each context category per cell line using Borzoi.

| Annotation | CRE type (deriveed from Enformer) | |
|---|---|---|
| | Enhancer | Silencer |
| Enhancer | 12.16 | 14.41 |
| TSS | 17.57 | 9.91 |
| Total | 27.03 | 23.87 |

**Supplementary Table 4.** Percentage overlap between sufficient enhancer and silencer CREs derived from Enformer versus annotations of enhancers (Enhancer Atlas), TSS (GENCODE) and combined annotations (Total).

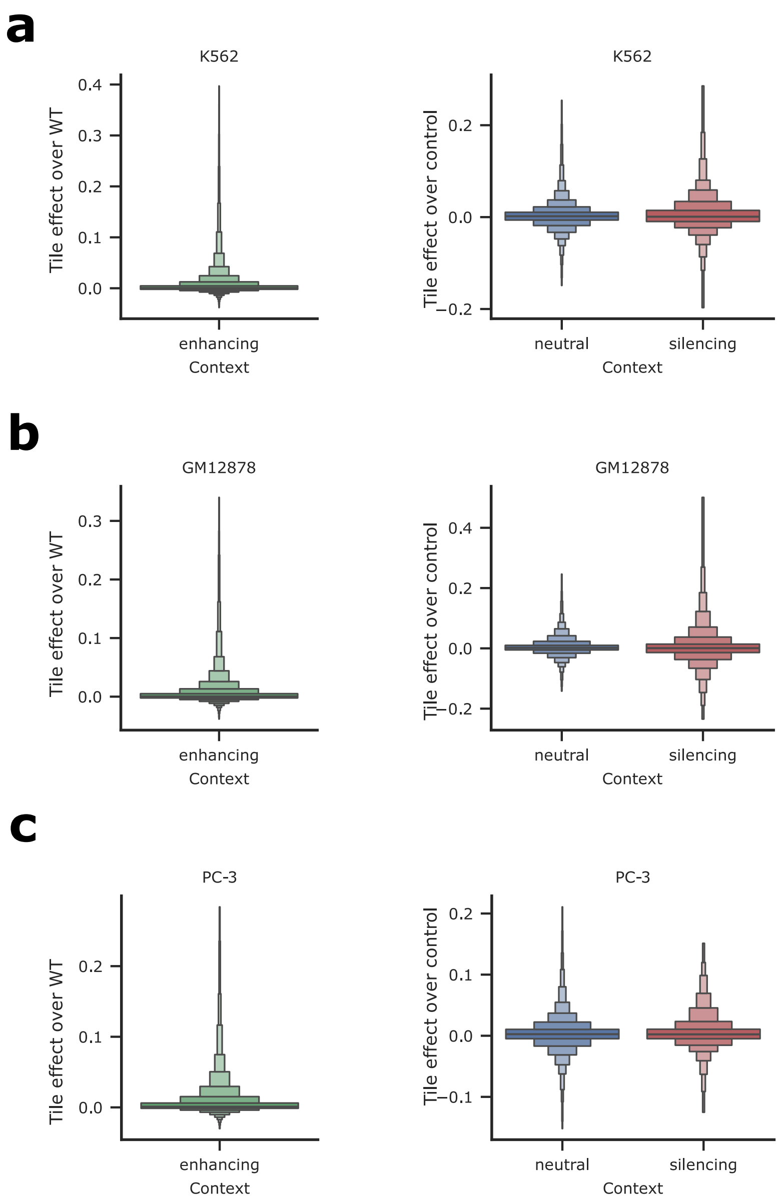


**Supplementary Figure 1.** Sufficiency test using Borzoi in three cell lines. Normalized CRE effect of embedding a given tile of interest along with the TSS tile into dinucleotide-shuffled sequences at their original positions for different context categories. The left panels in **a**-**c** show the results for enhancing context sequences normalized by intrinsic wild-type activity, and the right panels for neutral and silencing context sequences are normalized by intrinsic TSS activity. The number of data points in enhancing, neutral and silencing contexts is 20,400, 6,222 and 1,734 in panel **a**, 11,526, 5,508 and 1,224 in panel **b** and 7,242, 6,426 and 1,326 in panel **c**.

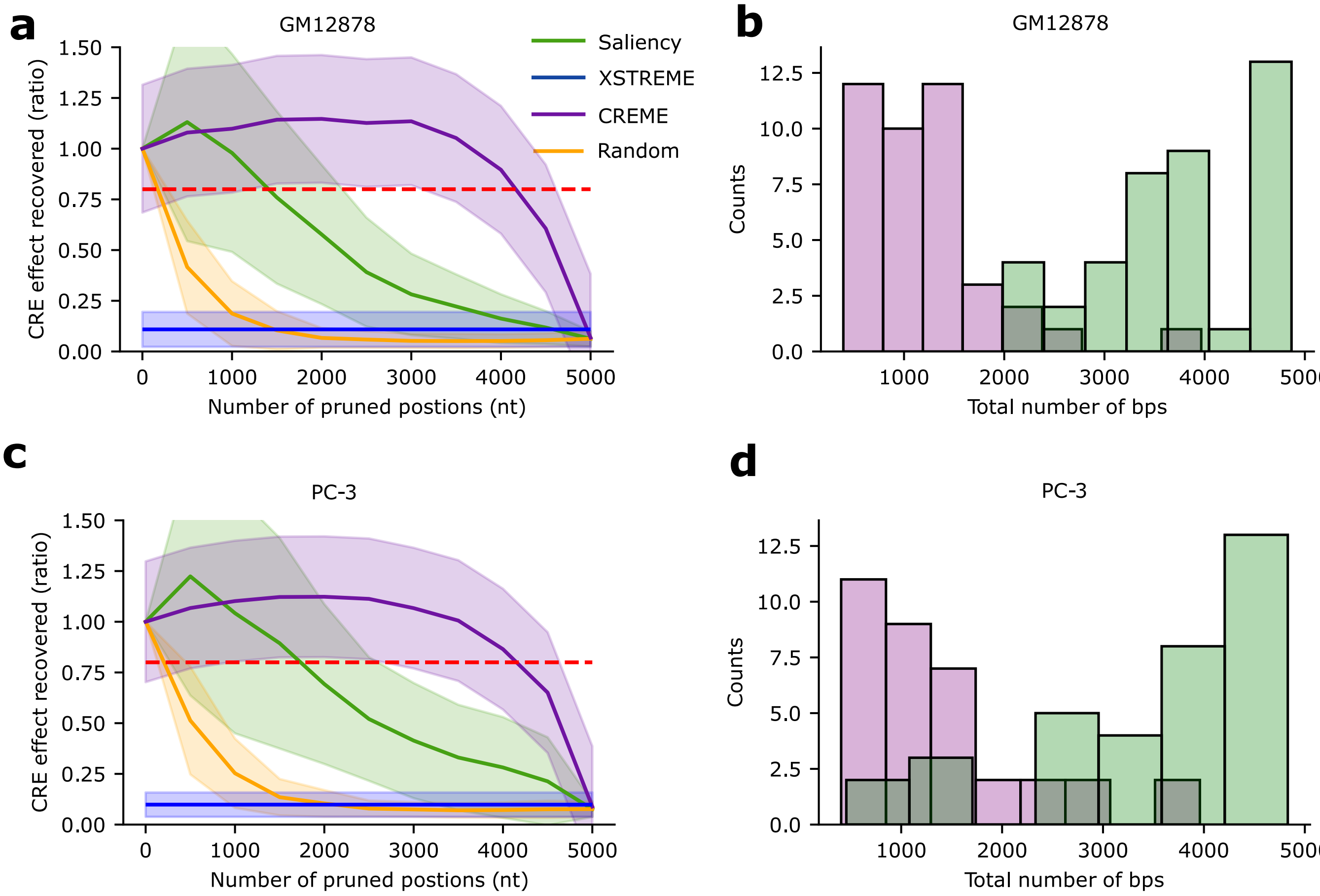


**Supplementary Figure 2.** Fine-tile search results for enhancing tiles in GM12878 and PC-3. **a**,**c**, Plot of the average CRE effect recovered ratio (which is the sufficiency of the sub-tiles divided by the sufficiency of the full tile) versus the number of pruned positions for enhancer tiles using different methods for GM12878 (**a**) and PC-3 (**c**). The red dotted line shows an arbitrary threshold of 0.8 (80%) explained. Shaded line represents the standard deviation of the mean. **b**,**d**, Histogram of number of remaining positions at a CRE effect recovered threshold of 0.8 for CREME versus saliency analysis for GM12878 (**b**) and PC-3 (**d**). The number of enhancer tiles explored is 41 and 35 for GM12878 and PC-3, respectively.

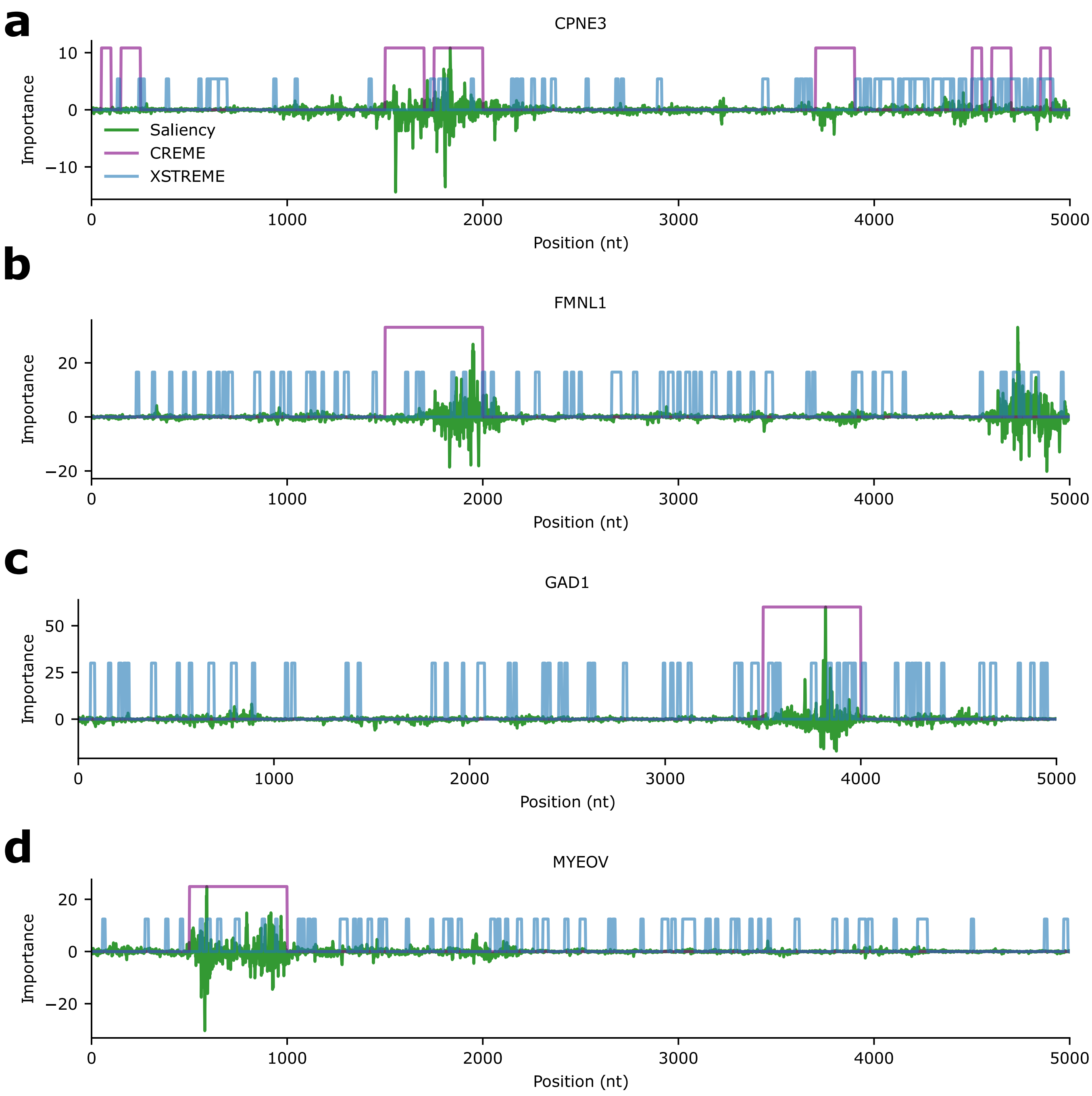


**Supplementary Figure 3.** Additional comparisons of sequence annotations within enhancer tiles for enhancer tiles in K562. **a**-**d**, Example annotations of important sequence elements for a putative enhancer for different genes.Each row shows an enhancer CRE of a different gene (indicated in the title). Examples of discrepancy (**a**,**b**) and agreement (**c**, **d**) between CREME and Saliency analysis. Note that the importance scale ($y$-axis) is set according to Saliency maps and the heights of binary annotations given by XSTREME and CREME are set arbitrarily for comparison.

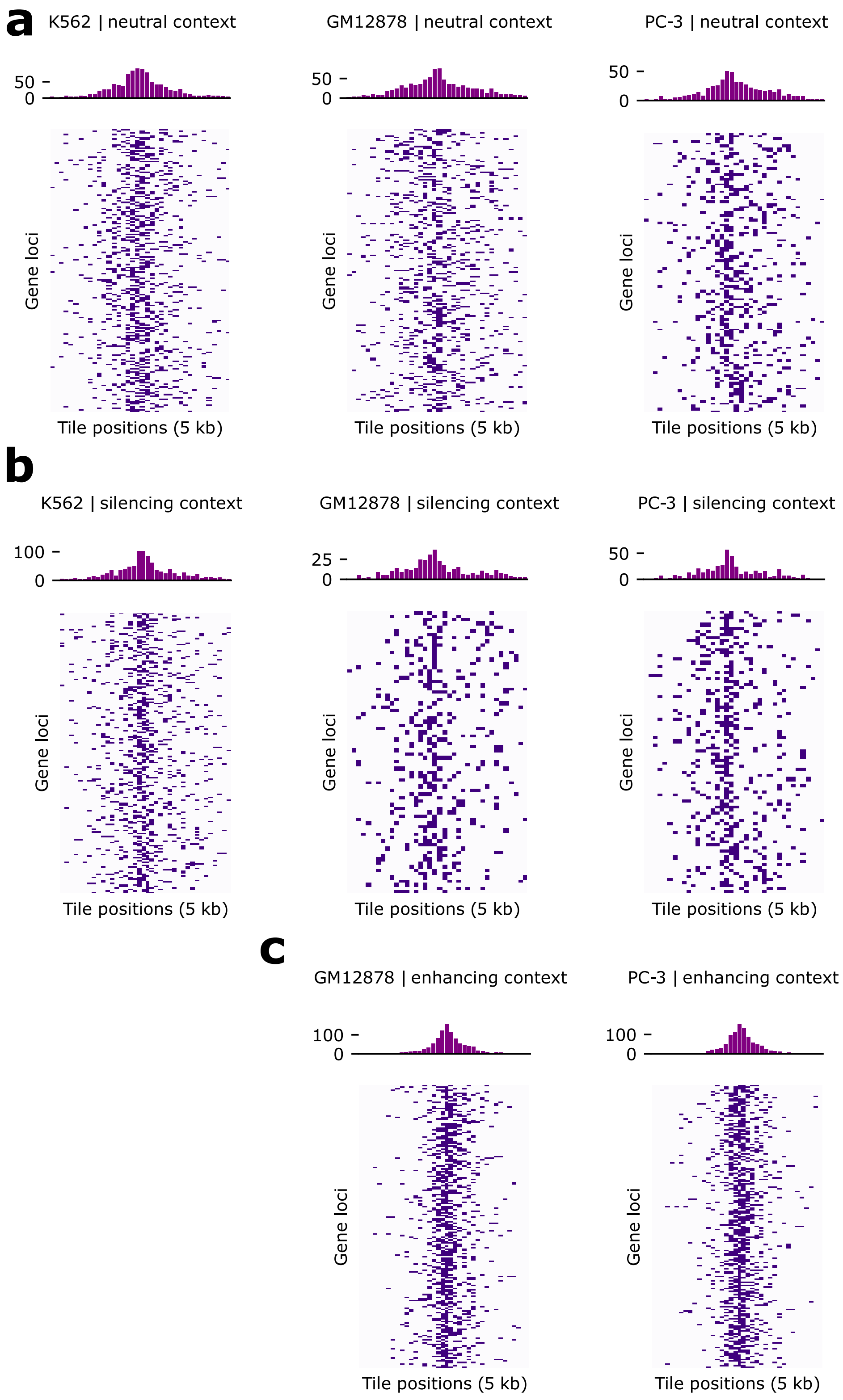


**Supplementary Figure 4.** Locations of identified CREs from the higher-order interaction test for enhancing elements across cell types using Enformer. **a**-**c**, Heatmaps show the locations of the first 5 CREs identified in the enhancer Higher-Order Interaction Test for sequences categorized as neutral, silencing or enhancing context, respectively. The histograms on top show the distribution of tile positions.

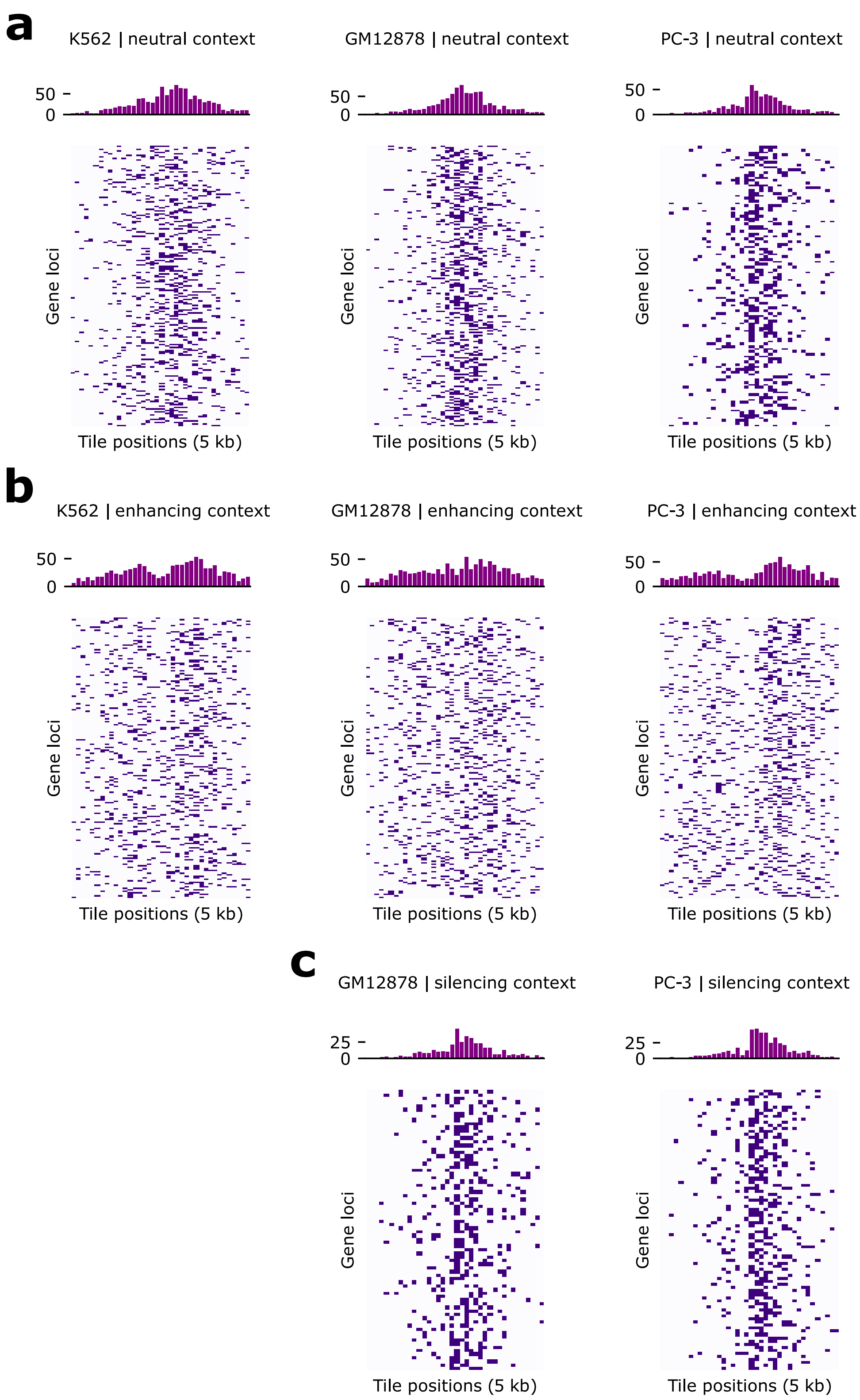


**Supplementary Figure 5.** Locations of identified CREs from the higher-order interaction test for silencing elements across cell types using Enformer. **a**-**c**, Heatmaps show the locations of the first 5 CREs identified in the silencer Higher-Order Interaction Test for sequences categorized as neutral, silencing or enhancing context, respectively. The histograms on top show the distribution of tile positions.

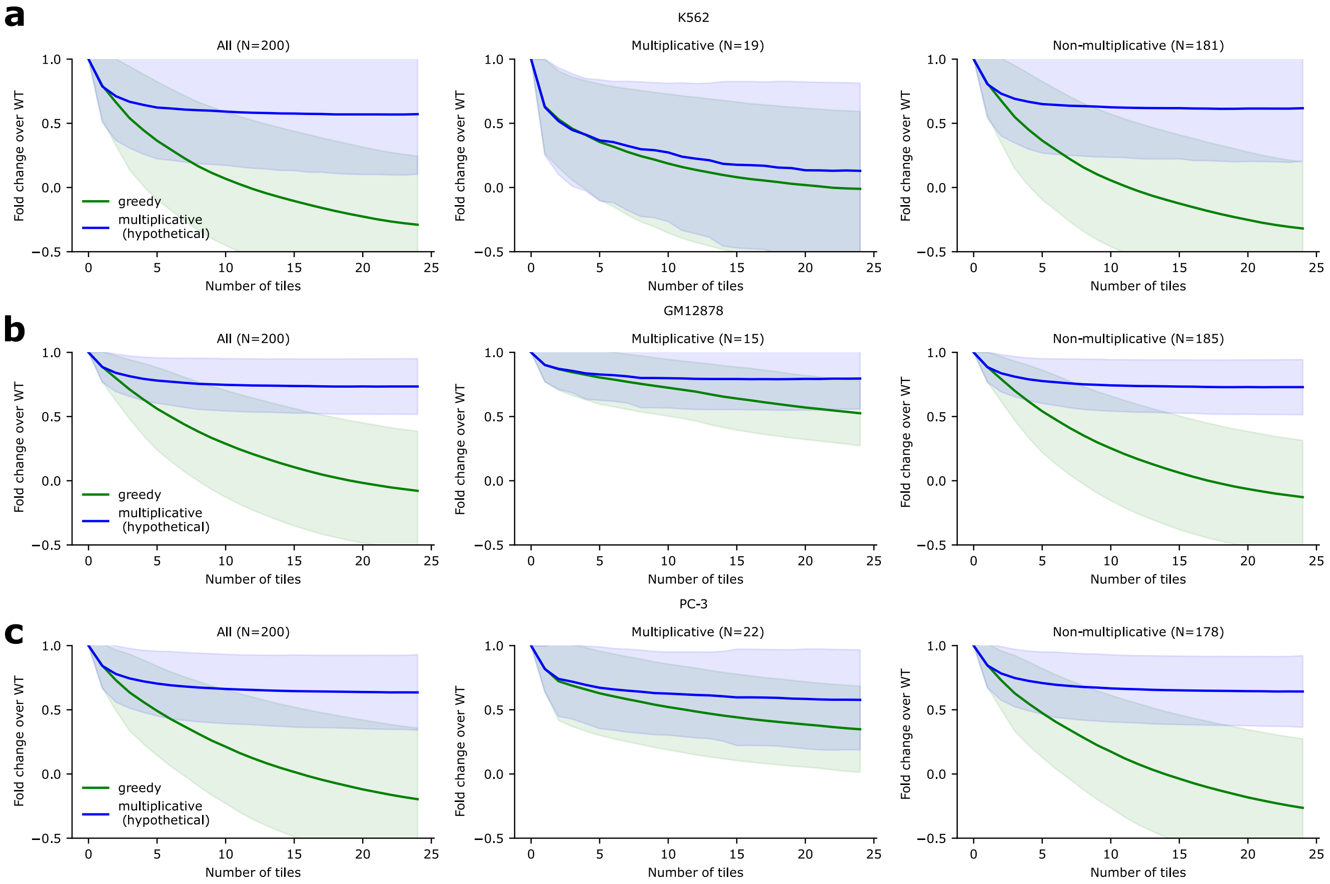


**Supplementary Figure 6.** Hypothetical multiplicative model of enhancer sets from the Higher-order Interaction Test. Comparison of the average fold change over WT for enhancer sets for sequences categorized as enhancing context versus a hypothetical multiplicative effects model. The 200 sequences from enhancing contexts are used in K562, GM12878 and PC-3 in panels **a**-**c**, respectively. Sequences are stratified into multiplicative or non-multiplicative according to the extent that they are concordant with the multiplicative model (see Methods).

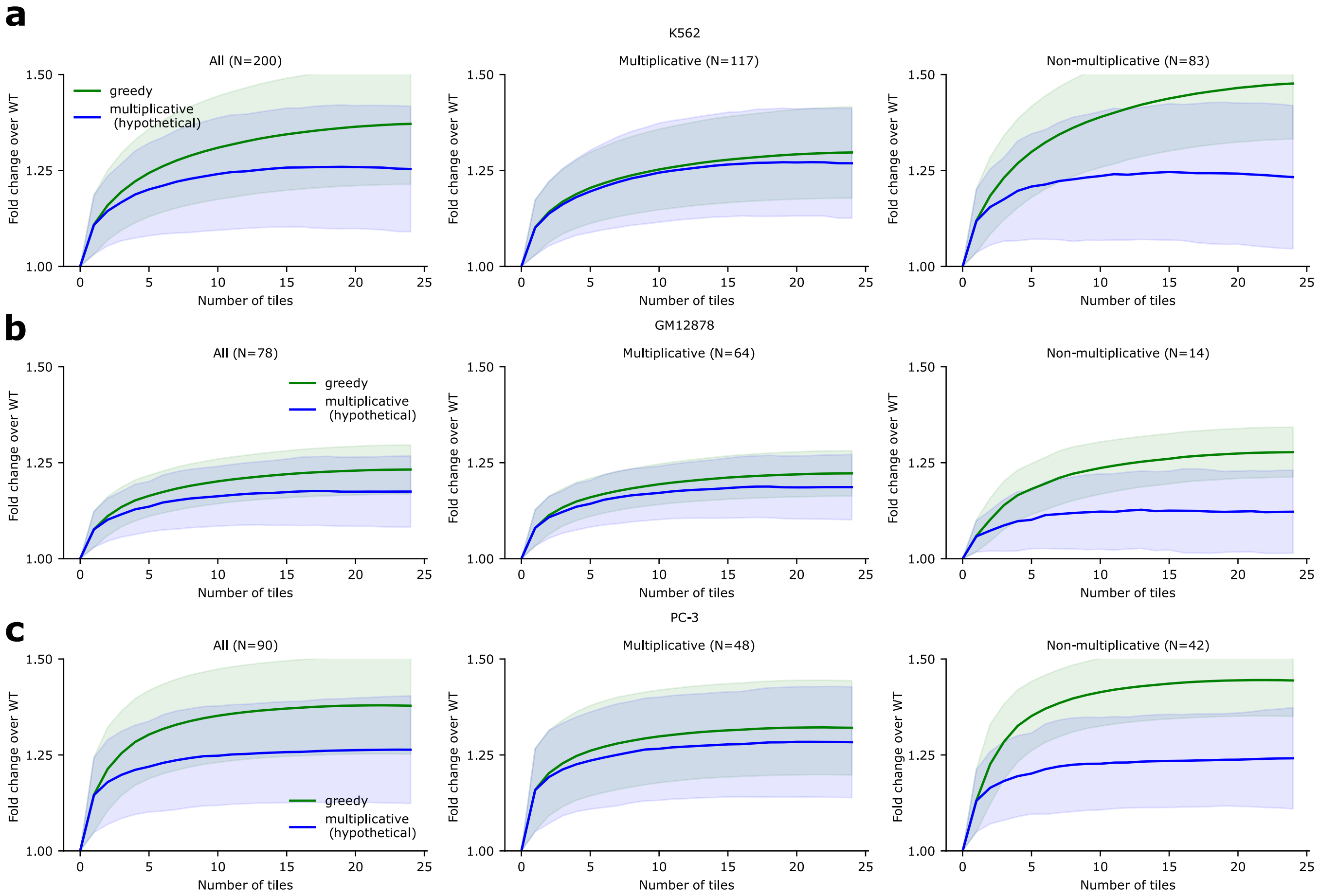


**Supplementary Figure 7.** Hypothetical multiplicative model of silencer sets from the Higher-order Interaction Test. Comparison of the average fold change over WT for silencer sets for sequences categorized as silencing context versus a hypothetical multiplicative effects model. The number of sequences from silencing contexts are shown for K562, GM12878 and PC-3 within the panels **a**-**c**, respectively. Sequences are stratified into multiplicative or non-multiplicative according to the extent that they are concordant with the multiplicative model (see Methods).
